# *Vibrio cholerae* interaction with predatory bacteria on chitin suggests an alternative mode of biofilm formation in marine snow conditions

**DOI:** 10.1101/2025.10.31.685833

**Authors:** Jacob D. Holt, Katherine A. Miller, Olivia F. Hunter, Emily Zhang, Alexander J. Hinbest, Emma Gerace, Rich Olson, Daniel E. Kadouri, Carey D. Nadell

**Affiliations:** Department of Biological Sciences, Dartmouth, Hanover, New Hampshire, USA; Department of Microbiology and Immunology, Geisel School of Medicine at Dartmouth, Hanover, New Hampshire, USA; Department of Oral Biology, Rutgers School of Dental Medicine, Newark, NJ, USA; Department of Molecular Biology and Biochemistry, Molecular Biophysics Program, Wesleyan University, Middletown, CT, USA

## Abstract

*Vibrio cholerae* is a ubiquitous marine microbe that solubilizes and consumes chitin in the marine water column. In both the marine environment and the intestinal track, *V. cholerae* forms biofilms; a key question regarding the lifestyle of *V. cholerae* is how do the diverse substrates that it encounters influence its biofilm formation and, in turn, shape its ecological interactions. Here, we use the predator-prey interaction between *Bdellovibrio bacteriovorus* and *V. cholerae* as a model to explore how the environmental chitin substrate alters *V. cholerae* biofilm formation and predator-prey interactions. We find that glass-bound biofilms provide strong protection for *V. cholerae* against predation while also allowing a population of predatory *B. bacteriovorus* to remain in place. In contrast, chitin-bound biofilms offer less protection against *B. bacteriovorus* predation and do not maintain a stable population of *B. bacteriovorus*. Using percolation and population dynamics models, we predict that these changes in predator-prey dynamics can be mostly explained by alterations in biofilm architecture between the two conditions, which changes the fraction of prey available to *B. bacteriovorus*. Performing targeted biofilm matrix deletions, we confirm this prediction by recapitulating key features of the chitin predator-prey interactions on glass surfaces. Following on this observation, we show that *V. cholerae* biofilms grown on chitin produce much less of the canonical biofilm matrix components and instead rely on other extracellular structures. Overall, our experiments detail how growth substrate can alter biofilm matrix composition and how these changes in biofilm architecture and cellular arrangement can impact higher-order ecological interactions.

## Introduction

*Vibrio cholerae* is the causative agent of cholera and a marine microbe that forms biofilms on the surface of chitin, where it secretes extracellular enzymes that degrade the insoluble polymer into soluble saccharides (1–5). Biofilm formation is a distinct physiological state that arises when planktonic cells become attached to a surface, reduce or halt motility, and secrete extracellular matrix composed of proteins, polysaccharides, and eDNA (6–8). Biofilm formation stabilizes bacterial colonization of nutrient-rich surfaces and in this instance links *V. cholerae* and other chitin-consuming marine bacteria to global carbon and nitrogen cycling (9–13). Additionally, biofilm formation provides protection against exogenous threats such as antibiotics, desiccation, bacteriophages, host immune systems, competing microbes, and predators - both prokaryotic and eukaryotic (14–24). While biofilm formation on glass surfaces has been well characterized in protecting *V. cholerae* from invading bacteria, bacteriophages, and predators, it remains unclear how environmental growth substrates of *V. cholerae*, such as chitin, may alter the protection offered by biofilm formation and what implications this may then have on biofilm community dynamics in aquatic environments (15–17, 23).

Recent work has indicated that the ecology and evolution of biofilm-dwelling bacteria are sensitive to the mechanistic details of matrix secretion and biofilm architecture (2, 24–27). For instance, for *V. cholerae* biofilms grown on glass under continuous media flow, disruption of the matrix protein RbmA leads to modified cell group architecture with greater spacing between cells and high sensitivity to invasion by competing strains, predatory bacteria, and phages (16, 23, 28, 29). Our goal in this paper was to utilize the predator-prey interaction between *Bdellovibrio bacteriovorus* and *V. cholerae* as a model to assess to what extent growth substrate alters biofilm architecture and predator-prey interactions under conditions recapitulating key features of the chitin marine snow surfaces that *V. cholerae* commonly occupies in the natural environment (3).

To assess the effect of chitin-bound growth on biofilm formation and predator-prey dynamics, we performed experiments simulating *V. cholerae* biofilm growth on chitin particles and exposed them to *B. bacteriovorus*, an obligate predator of Gram negative bacteria (30). We also conducted comparison experiments on glass surfaces using one of two different carbon sources: the soluble disaccharide (GlcNAc)_2_ or the soluble monosaccharide GlcNAc. the primary end-products of *Vibrio*-secreted chitinases (4). We then employed quantitative biofilm architecture measurements, ecological and physical models, and genetic manipulation of matrix production and composition to assess how growth on chitin changes *V. cholerae* biofilm architecture, and how these changes in turn impact the biofilm population’s interaction with predatory bacteria. Altogether, our findings show that the details of surface association and growth substrate can feed back to biofilm matrix regulation and composition, and how these changes in turn can cascade to influence higher order ecological interactions.

## Results

### *V. cholerae* biofilms formed on chitin are more susceptible to predation by *B. bacteriovorus* than biofilms formed on glass

To observe live-cell population dynamics in biofilms under fluid flow, we used strains of *V. cholerae* and *B. bacteriovorus* that had been engineered to constitutively express the fluorescent proteins mKate2 and GFP, respectively, allowing them to be distinguished by live-cell fluorescence microscopy. *V. cholerae* was inoculated into microfluidic devices composed of PDMS chambers bonded to glass coverslips; the chambers contained V-shaped traps that were pre-loaded with chitin flakes to which *V. cholerae* cells could adhere (SI Figure 1). After a 1 h attachment period, *V. cholerae* was incubated under continuous flow of M9 minimal medium – with either no carbon source (for experiments in which biofilms grew on chitin), (GlcNAc)_2_, or GlcNAc – at 0.1 µL/min (average flow velocity ∼ 14 µm/s). After 72 h of *V. cholerae* biofilm formation, *B. bacteriovorus* was inoculated into the existing *V. cholerae* biofilm chambers via a tubing swap to media containing 10^10^ PFU/mL of *B. bacteriovorus*. After a 2 h invasion period, the tubing was swapped back to the original media, and imaging was performed 2 h later and every 24 h thereafter until the end of the experiment.

**Figure 1:**
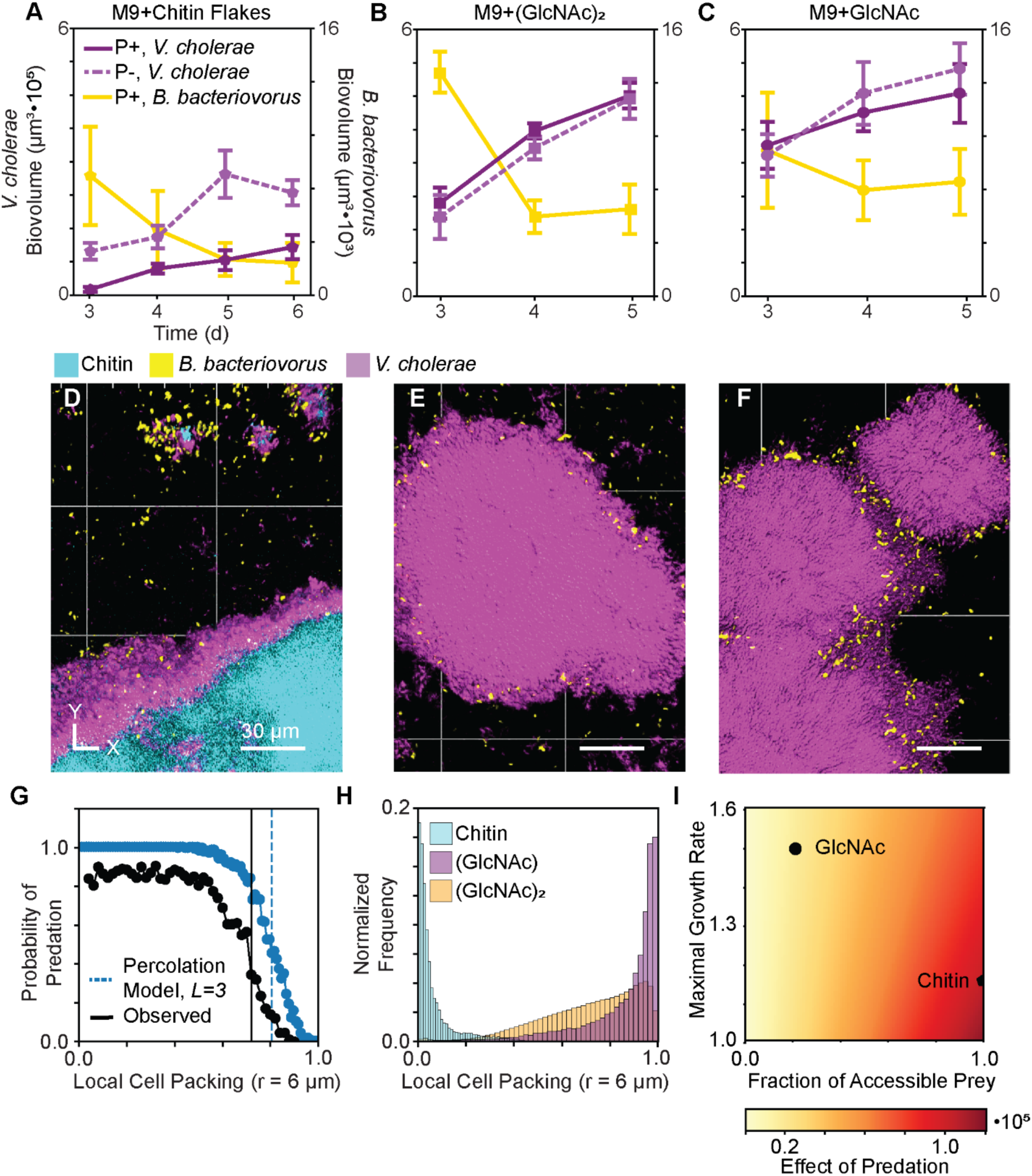
*V. cholerae* biofilm populations on chitin are more susceptible to *B. bacteriovorus* predation than biofilms formed on glass with GlcNAc or (GlacNAc)_2_ as the carbon source. **(A)** Population dynamics of *V. cholerae* biofilms formed on chitin with or without *B. bacteriovorus* introduced after 3 d of growth (P+ solid line and P– dashed line, respectively). **(B)** Population dynamics of *V. cholerae* biofilms formed on glass with (GlcNAc)_2_ in the medium, with or without the addition of *B. bacteriovorus*. **(C)** Population dynamics of *V. cholerae* biofilms formed on glass with GlcNAc in the medium, with or without the addition of *B. bacteriovorus*. **(D)** Representative 3D projection of a *V. cholerae* biofilm on chitin, following addition of *B. bacteriovorus*. **(E)** Representative 3D projection of a (GlcNAc)_2_ supplied *V. cholerae* biofilm on glass, following addition of *B. bacteriovorus*. **(F)** Representative 3D projection of a GlcNAc-supplied *V. cholerae* biofilm on glass, following addition of *B. bacteriovorus*. For all image panels, *V. cholerae* is shown in purple, *B. bacteriovorus* is shown in yellow, and chitin is shown in cyan. **(G)** Probability of predation as a function of local cell packing plotted from experimental data and from simulation results. **(H)** Histograms of WT *V. cholerae* biofilm local cell packing across three media conditions: M9+GlcNAc, M9+(GlcNAc)_2_, and M9+Chitin. **(I)** Heatmap showing how the effect of the predator on prey abundance, defined as the time-average difference between the predator treated and untreated prey populations, changes as the maximal growth rate is varied from 0.5 to 1.5 and the fraction of accessible prey is varied from 0 to 1. The chitin and GlcNAc model fits are mapped to the heatmaps as a pentagon and circle.

When grown in M9 minimal media with chitin as the sole carbon source, a significant decrease in the biovolume of *B. bacteriovorus*-exposed biofilms was observed relative to control biofilms that received a tubing swap with sterile media (Fig 1A, SI Figure S2A). We next sought to determine if *V. cholerae* biofilms grown on chitin are more or less susceptible to predation by *B. bacteriovorus* relative to biofilms grown on glass with nutrients supplied in the surrounding liquid; to do this we performed experiments in which *V. cholerae* was grown on the glass substrate of the chambers with one of either GlcNAc or (GlcNAc)_2_ (0.5%) as the sole carbon source in the liquid media (31). In contrast to the chitin condition, when *V. cholerae* was grown in M9 media on glass with either GlcNAc or (GlcNAc)_2_ as the sole carbon source, its biofilms were much more resilient to *B. bacteriovorus* exposure. Predation events could be observed along the biofilm periphery, and *B. bacteriovorus* remained associated with the host biofilm, but there was no significant net effect of predation on *V. cholerae* population dynamics (Fig 1B,C SI Figure S2B,C).

From this initial set of experiments, we noted three main features distinguishing *V. cholerae* biofilms formed on chitin particles versus biofilms grown on glass with GlcNAc or (GlcNAc)_2_ in the media. First, the total abundance of the *V. cholerae* growing on chitin was substantially lower than for biofilms on glass with growth substrate in the influent media (Fig 1A-C, SI Figure S2). Second, following the introduction of *B. bacteriovorus* and initial bouts of predation, the total abundance of *B. bacteriovorus* remaining biofilm-associated appeared to continually drop within biofilms grown on chitin, while *B. bacteriovorus* abundance quickly stabilized within biofilms grown in GlcNAc or (GlcNAc)_2_. Third, *V. cholerae* biofilms grown on chitin appeared to have lower internal cell packing and to allow *B. bacteriovorus* to permeate most of their interior volume, while biofilms grown on glass appeared to be more densely packed with cells and confined *B. bacteriovorus* to the outside of mature cell groups (Fig 1D-F, SI Figure S2). The third observation recapitulates our prior findings, which found a consistent, negative relationship between the volume fraction occupied by prey bacterial cells within biofilms and the likelihood that a given cell would be predated by *B. bacteriovorus* (16, 27).

We next assessed whether a simple percolation model that considers only the occupation of space by prey cells could predict the likelihood that a focal prey cell could be reached by *B. bacteriovorus*. The percolation model gave a good match to measurements of predation probability as a function of nearby cell packing across our experiments (Fig 1G, SI Figure S3). This result in turn suggested that the difference in predator-prey population dynamics between biofilms grown on chitin versus glass could be largely due to differences in prey cell accessibility based on cell packing within the biofilm. To test this possibility we used a modified Lotka-Volterra model to ask whether the observed difference in cell packing between *V. cholerae* biofilms grown on chitin versus glass (Fig 1H) – and the resulting predicted difference in predator exposure – could explain the observed difference in predator-prey population dynamics between the different growth conditions. This approach produced a partial match to the experimental data, connecting the difference in cell packing observed between the chitin- and glass-grown biofilm conditions and the observed impact of *B. bacteriovorus* exposure on prey abundance over time (Fig 1I, SI Figure S4).

However, our model did not capture the dynamics of *B. bacteriovorus*. We were not able to find a case in which the predator maintained stable positive abundance while having a lower effect on prey abundance; instead, we observed a rapid decline of predator abundance in all model cases (SI Figure S5). Further exploring our model, we determined that to see increased predator abundance as the predator’s available prey pool decreases, there must be a compensatory effect where either the predator’s loss rate decreases with decreasing prey accessibility, or the predator’s prey conversion efficiency increases with decreasing prey accessibility (SI Figure S5). This pointed towards an additional subtlety in the experiments, beyond simply where predators and prey cells are located, which we explore further below. Altogether these results suggest fundamental differences in *V. cholerae* biofilm structure on chitin versus on glass under flow of GlcNAc or (GlcNAc)_2_. In the following section, we manipulate *V. cholerae* matrix composition on glass surfaces to test the hypothesis that relatively simple changes in matrix composition could account for the major differences in predator-prey interaction on chitin versus glass that we have documented thus far.

### Partially matrix-deficient biofilms formed on glass recapitulate key features of *B. bacteriovorus*-*V. cholerae* interaction on chitin

To further define the relationship between matrix composition, cell arrangement, and predator-prey population dynamics, we performed experiments using strains harboring deletions of key extracellular matrix components grown on glass, focusing on RbmA and the core *Vibrio* exopolysaccharide (VPS). RbmA is a secreted protein that interacts with the *V. cholerae* cell surface and other matrix components and plays an important role in controlling how closely cells are arranged together (28, 32–34). Mutants lacking RbmA can still produce biofilms, but with substantially reduced cell packing (Fig 2A,B) (8, 28). While we previously showed that deletion of *rbmA* leads to high predator invasion and susceptibility in comparison to WT *V. cholerae* biofilms, the goal here was to determine the degree to which targeted matrix deletions in biofilms formed on glass could recapitulate key features of *B. bacterivorous* predation of WT biofilms formed on chitin (16, 27).

**Figure 2:**
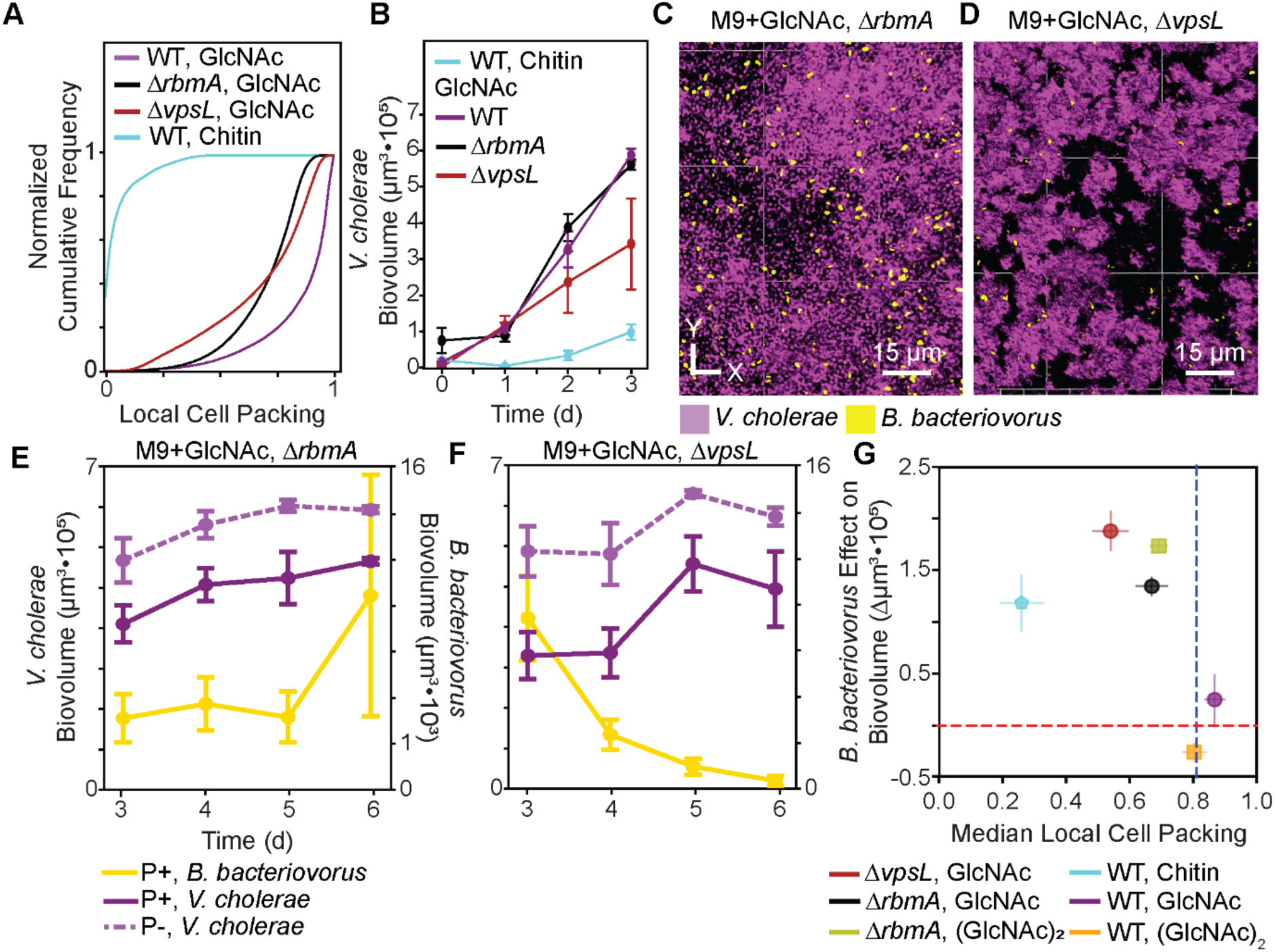
Abolishment of *Vibrio* exopolysaccharide (VPS) secretion recapitulates features of chitin predator-prey dynamics on glass substrate. **(A)** Cumulative frequency histogram of local cell packing for WT grown on chitin, WT grown in GlcNAc, *ΔvpsL* grown in GlcNAc, and *ΔrbmA* grown in GlcNAc. **(B)** Biofilm growth curves for WT grown on chitin, WT grown in GlcNAc, *ΔvpsL* grown in GlcNAc, and *ΔrbmA* grown in GlcNAc. **(C)** Representative image of a *ΔrbmA* biofilm being predated upon by *B. bacteriovorus*. **(D)** Representative image of a *ΔvpsL* biofilm being predated upon by *B. bacteriovorus*. *V. cholerae* is shown in purple, *B. bacteriovorus* is shown in yellow, and chitin is shown in cyan. **(E)** Population dynamics of *ΔrbmA V. cholerae* biofilms formed on chitin with *B. bacteriovorus* added after 3 d of growth (P+, solid line) and without the addition of *B. bacteriovorus* (P–, dashed line). **(F)** Population dynamics of *ΔvpsL V. cholerae* biofilms formed on chitin with *B. bacteriovorus* added after 3 d of growth and without the addition of *B. bacteriovorus.* **(G)** The effect of *B. bacteriovorus* on *V. cholerae* biofilm biovolume plotted against the median monoculture cell packing for five conditions: WT grown on chitin shown in cyan, WT grown in GlcNAc shown in purple, WT grown in (GlcNAc)_2_ shown in orange, *ΔvpsL* grown in GlcNAc shown in red, and *ΔrbmA* grown in GlcNAc shown black. The critical threshold, *p_c_*, predicted by the percolation model in Figure 1G is shown as a dashed blue line.

Repeating the predation experiments from Figure 1 with biofilms formed by the *ΔrbmA* strain, we recapitulated the result that *B. bacteriovorus* permeates through most of the biofilm, homogeneously spreading out within the host population, rather than being restricted to the outer periphery (16, 27). Additionally, as in the chitin case, we observed a reduction in *V. cholerae* abundance compared to a no-predator control (Fig 2E, SI Figure S6). However, unlike in the chitin case, *B. bacteriovorus* is retained within *ΔrbmA* biofilms at an abundance similar to WT biofilms grown on glass substrate. (Fig 1B,C, Fig 2E, SI Figure S6). While the loss of RbmA within biofilms on glass recapitulated the increased susceptibility of those formed on chitin, it was unable to account for the decrease in predator abundance observed in the chitin experiments. This result motivated us to turn to another biofilm matrix component that also results in decreased local cell packing on glass, VPS (Fig 2A).

To characterize the interaction of *B. bacteriovorus* with *V. cholerae* biofilms producing substantially decreased VPS amounts, we took advantage of a previously characterized *vpsL* deletion mutant. VpsL is a key component of the matrix exopolysaccharide secretion machinery, and mutants lacking VpsL do not secrete VPS (33). Because the matrix proteins RbmA, RbmC, and Bap1 all directly interact with VPS, biofilms lacking VPS are thought not to retain the three primary matrix proteins, either. Consequently, *ΔvpsL* mutants either fail to produce 3-dimensional biofilms or form irregularly shaped biofilms with varying degrees of 3-dimensional structure, dependent on media conditions (SI Figure S7) (2, 33, 35–40). In M9+GlcNAc, *V. cholerae ΔvpsL* produces irregularly shaped 3-dimensional biofilm clusters with cell packing profiles that are reduced compared to VPS-producing cell groups (Fig 2A,B,D).

Repeating the predation experiment with *V. cholerae ΔvpsL* grown on glass with GlcNAc, we observed *B. bacteriovorus* initially permeating most of the *V. cholerae* population and a significant decline in *V. cholerae* abundance compared to a no-predator control (Fig 2D,F, SI Figure S6). While *V. cholerae* biofilm susceptibility to predation is similar between the *ΔvpsL* and the *ΔrbmA* mutants under these culture conditions, the effects of each deletion mutant on *B. bacteriovorus* population dynamics are different. Though they are able to attack and consume prey cells within 24 h of being introduced, *B. bacteriovorus* did not persist within or along the periphery of *ΔvpsL* biofilms in the days following (Fig2F, SI Figure S6). Altogether these results indicated that the ability of *B. bacteriovorus* to be maintained on *V. cholerae* biofilms grown on glass substrate is dependent on the presence of VPS and potentially its interactions with other matrix components, and not solely on the abundance of accessible prey.

So far, we have confirmed that, regardless of underlying surface, lowered local cell packing results in an increased susceptibility to predation by *B. bacteriovorus* (Fig 2G). However, only the inhibition of VPS secretion is sufficient to recapitulate both the increased susceptibility to predation and the decreased predator survival observed previously in the chitin experiments. These results suggest that biofilms formed on chitin may not be producing VPS, or may produce it in much lower quantities.

### Biofilms formed on chitin have altered matrix configuration relative to biofilms formed on glass

The observations above prompted use to ask whether VPS and the other primary matrix components, RbmA, Bap1, and RbmC were being produced in lower quantities when *V. cholerae* is on chitin (25, 27). To assess production of VPS we used an approach described by Huang et al. (2023) in which the β-propeller domain of Bap1, which binds to VPS, is conjugated to a fluorophore to serve as a VPS-specific fluorescent stain (36). We observed diffuse and bright stain signal through biofilms grown in GlcNAc on glass, while biofilms formed on chitin showed little to no signal (Fig 3A, SI Figure S8A). Using a previously described immunostaining approach for the secreted proteins RbmA, Bap1, and RbmC, we also observed little matrix protein production within biofilms grown on chitin relative to biofilms formed on glass with GlcNAc in the surrounding medium (Fig 3B,C,D, SI Figure S8B,C,D) (16, 23, 25, 34).

**Figure 3:**
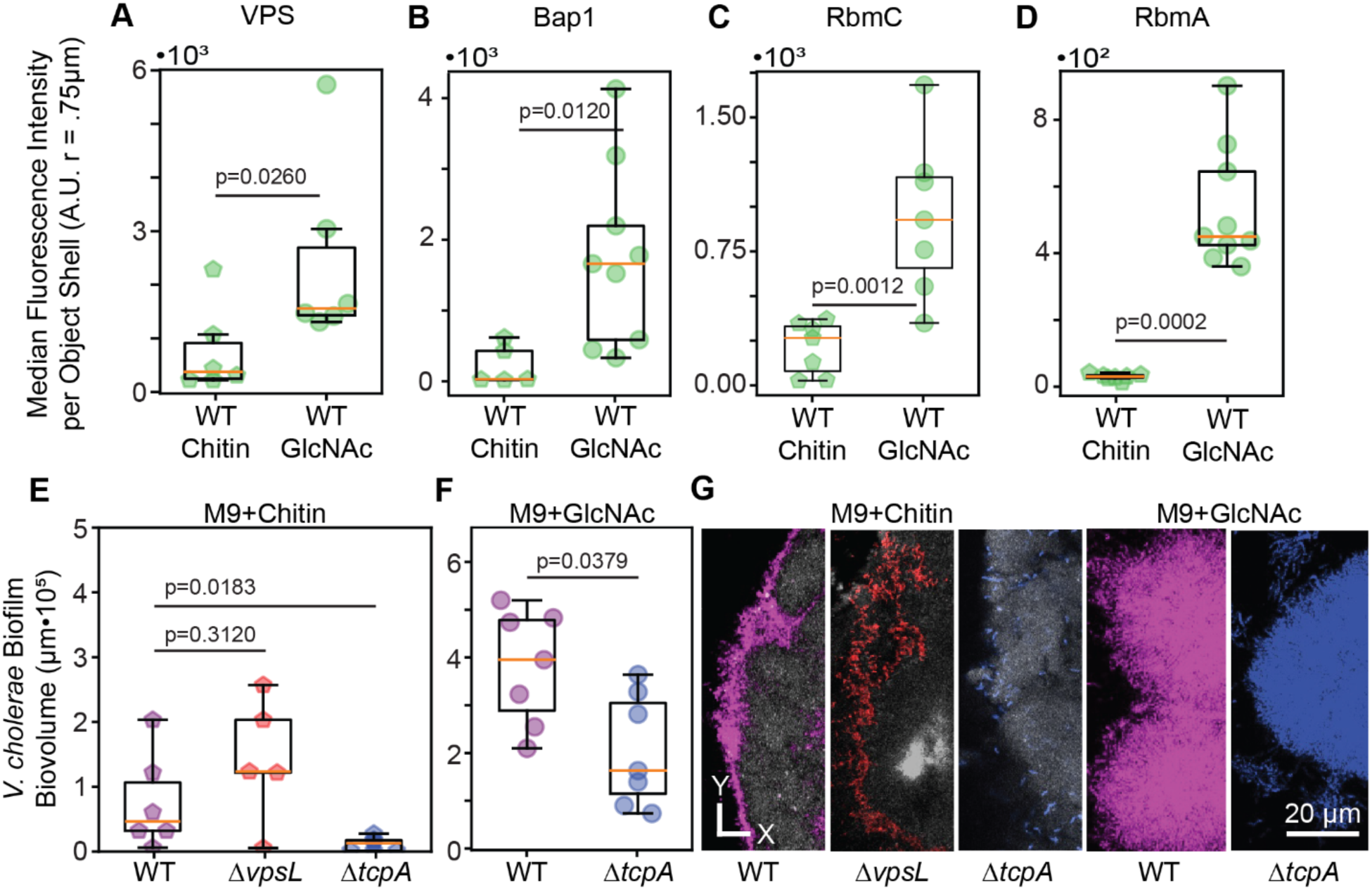
Growth on chitin shifts biofilm formation from VPS-dependent to TCP-dependent. **(A)** Staining of VPS quantified as the median pseudo-cell cube fluorescence intensity within a 0.75 μm shell is lower on chitin relative to GlcNAc (Mann-Whitney *U* test, n=6, p=0.0260). **(B)** Immunostaining of FLAG tagged Bap1 quantified as the median pseudo-cell cube fluorescence intensity within a 0.75 μm shell is significantly less on chitin relative to GlcNAc (Mann-Whitney *U* test, n=5-9, p =0.0120). **(C)** Immunostaining of FLAG tagged RbmC quantified as the median pseudo-cell cube fluorescence intensity within a 0.75 μm shell is significantly less on chitin relative to GlcNAc (Mann-Whitney *U* test, n=7, p=0.0012). **(D)** Immunostaining of FLAG tagged RbmA quantified as the median pseudo-cell cube fluorescence intensity within a 0.75 μm shell is significantly less on chitin relative to GlcNAc (Mann-Whitney *U* test, n=7-9, p=0.0002). **(E)** Biofilm volume at 3 d of *V. cholerae* WT, *ΔvpsL*, and *ΔtcpA* grown with chitin as the sole carbon source showing a significant decrease between WT and *ΔtcpA* (Mann-Whitney *U* test, n=6-7, p=0.0183) and no significant difference between WT and *ΔvpsL* (Mann-Whitney *U* test, n=5-6, p=0.3120). WT data replotted from Fig. 2. **(F)** Biofilm volume at 3 d of *V. cholerae* WT and *ΔtcpA* grown with GlcNAc as the sole carbon source showing a significant decrease (Mann-Whitney *U* test, n=7, p=0.0286). **(G)** Representative images of WT, *ΔvpsL*, and *ΔtcpA* growing on chitin and GlcNAc. WT is shown in purple, *ΔvpsL* is shown in red, *ΔtcpA* is shown in blue, and chitin is shown in gray.

From the results above we inferred that for our strains and culture conditions, VPS-dependent biofilm formation as a whole is activated to a much lower degree when *V. cholerae* is growing on chitin. This interpretation predicts that a *ΔvpsL* mutant, which secretes little or no VPS, should grow similarly to its isogenic parental strain on chitin, which we confirmed (Fig 3E,G). Further supporting our interpretation of decreased VPS-dependent biofilm production on chitin, in two-strain competition experiments performed on chitin or on glass with GlcNAc, the *ΔvpsL* mutant had a higher fitness against its parental strain on chitin relative to the glass and GlcNAc growth condition (SI Figure S9A,B). These outcomes, together with the matrix component staining results above, prompted us to consider other possibilities for cellular structures holding cells to each other in association with the chitin surface in these experiments.

Prior studies have suggested that the Type 4 pili of *V. cholerae* can be used for aggregation, as well as biofilm formation in some contexts (37, 41–46). Specifically, in our strain background, N16961, the Toxin Co-regulated Pilus (TCP) has been implicated as an important structure for biofilm integrity on chitin derived from squid pens (47). To test if the TCP is related to biofilm formation in our experiments, we constructed a *ΔtcpA* deletion mutant, abolishing TCP biosynthesis, and assessed the impact on chitin-bound biofilm growth.

After 3 d of incubation on chitin, *ΔtcpA* biofilms exhibited significantly reduced accumulation compared to the WT control (Fig 3E,G). For comparison, we repeated this experiment using glass as the substratum for biofilm growth and GlcNac as the sole carbon source in the surrounding medium. In this context, the *ΔtcpA* strain also showed reduced biomass compared to WT, but it was able to produce microcolony structures indicative of well-studied VPS-dependent biofilm formation (Fig 3F,G). The *ΔtcpA* mutant also had a competitive disadvantage against WT in biofilms on glass with GlcNAc and on chitin, further supporting a role for TCP in biofilm integrity (SI Figure S9C,D). These results align with prior work and suggest that biofilm formation on chitin is dependent on production of one or more pili (41, 47). While it was surprising to us that the *ΔtcpA* strain showed a phenotype on glass surfaces, this result is consistent with reports that *tcpA* expression levels are elevated in biofilms formed both on and off of chitin (48).

### A hyperactive diguanylate cyclase mutant forms biofilms on chitin that are protected from *B. bacteriovorus* predation

If our inference that VPS-dependent biofilm growth is reduced on chitin in favor of other mechanisms that could hold cells together in proximity to the surface, we predict that increasing VPS-dependent biofilm growth in the chitin environment via genetic modification will restore a similar degree of prey protection and predator abundance as seen in biofilms grown on glass. To test this, we used a strain of *V. cholerae* harboring a point mutation, *vpvC^W240R^*, previously characterized as elevating intracellular c-di-GMP concentration and leading to constitutive production of VPS and the matrix proteins RbmA, RbmC, and Bap1 (16, 49). We confirmed this strain behaves as expected in our culture conditions using RbmA immunostaining (Fig 4A-D, SI Figure S11). RbmA immunostaining correspond strongly with VPS staining, making it suitable as a general matrix stain in *V. cholerae* biofilms (SI Figure S10) (16).

**Figure 4:**
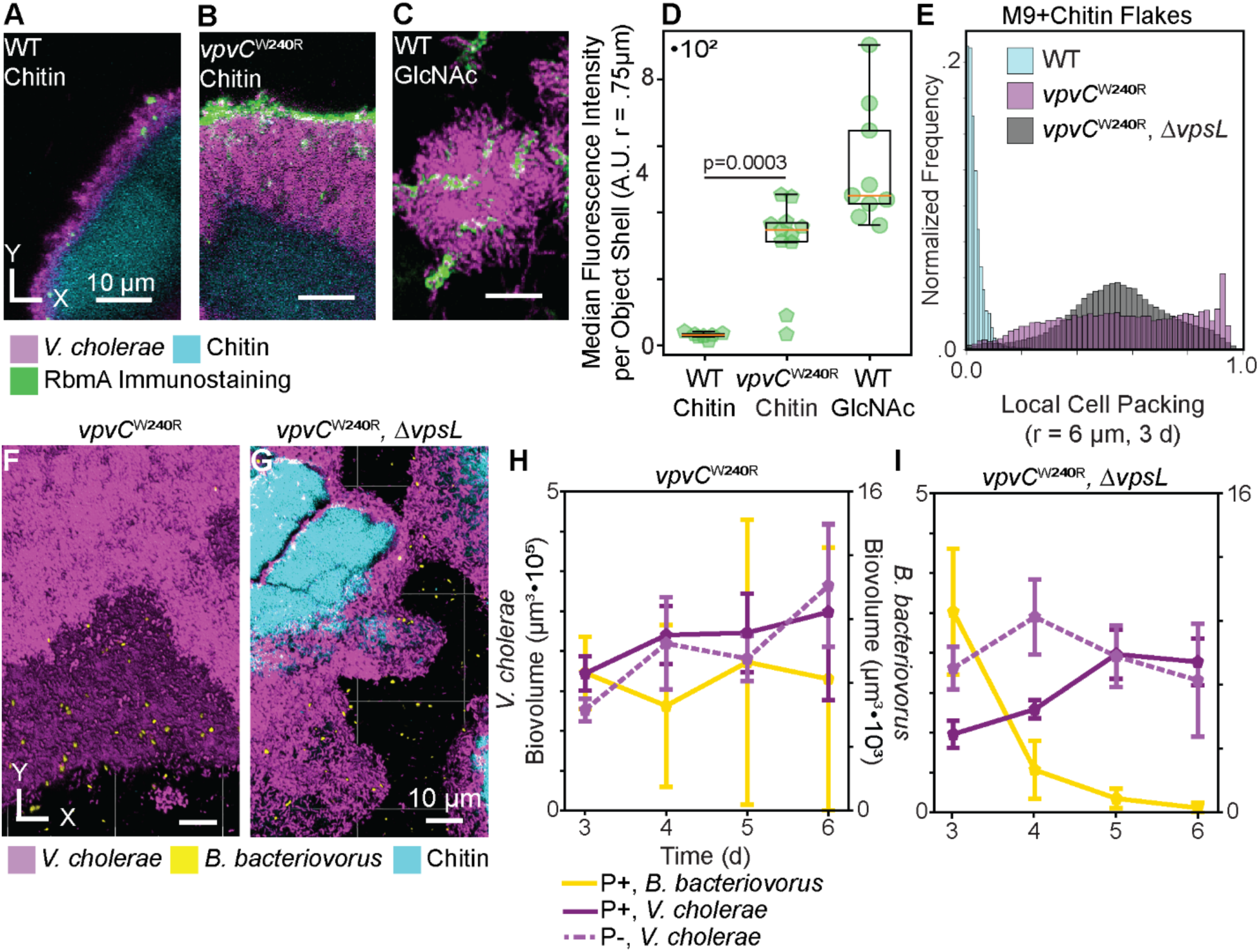
A hyperactive diguanylate cyclase mutant, *vpvC^W240R^*, forces VPS-dependent biofilm formation on chitin, resulting in predator-prey dynamics to what is observed in biofilms on glass. **(A)** Representative image of chitin grown WT RbmA immunostaining. **(B)** Representative image of chitin grown *vpvC^W240R^* RbmA immunostaining. **(C)** Representative image of GlcNAc grown WT RbmA immunostaining. *V. cholerae* is shown in purple, RbmA immunostaining signal is shown in green, and chitin is shown in cyan. **(D)** Box and whisker plot showing a significant increase in RbmA immunostaining between *vpvC^W240R^* and WT on chitin (Mann-Whitney *U* test, n=7-11, p=0.0036, WT data replotted from Fig. 3). **(E)** Cell packing frequency distributions of WT, *vpvC^W240R^*, and *vpvC^W240R^*, *ΔvpsL* biofilms grown on chitin. **(F)** Representative 3D rendering of a *vpvC^W240R^* biofilm (purple) being predated upon by *B. bacteriovorus* (yellow). **(G)** Representative 3D rendering of a *vpvC^W240R^, ΔvpsL* biofilm being predated upon by *B. bacteriovorus*. Chitin is shown in cyan. **(H)** Population dynamics of *vpvC^W240R^* with (P+, solid line) and without (P–, dashed line) predation. **(I)** Population dynamics of *vpvC^W240R^, ΔvpsL* with and without predation. *V. cholerae* biofilm without predator addition is shown as a dashed purple line. *V. cholerae* biofilm with predator addition is shown as a solid purple line. *B. bacteriovorus* is shown as a yellow solid line.

*vpvC^W240R^* biofilms grown on chitin have substantially increased cell packing relative to a WT control. Because the *vpvC^W240R^*mutation increases c-di-GMP levels, which can influence the expression of many loci, it was also important to show that changes in *vpvC^W240R^*biofilm morphology on chitin were specifically due to VPS-dependent matrix, rather than other factors. We performed this check by making a *vpsL* deletion in the *vpvC^W240R^* background, which resulted in decreased cell packing relative to *vpvC^W240R^* when grown on chitin. The double mutant strain did still produce more tightly packed biofilms than the WT on chitin, however, suggesting moderate roles for other factors altered by elevated c-di-GMP pools (Fig 4E) that we do not explore further here. Invading *B. bacteriovorus* into mature *vpvC^W240R^* biofilms formed on chitin, we observed *B. bacteriovorus* localization to the biofilm exterior, a negligible effect of *B. bacteriovorus* on *vpvC^W240R^*biofilm abundance over time, and increased retention of *B. bacteriovorus*, consistent with our predictions and data from the previous sections (Fig 4F,H, SI Figure S11). Consistent with the prior experiments with biofilms lacking VPS, *B. bacteriovorus* has a measurable effect on the abundance of *vpvC^W240R^*Δ*vpsL* biofilms at early timepoints, but as *B. bacteriovorus* is lost over time the effect of the predator is no longer observed (Fig 4G,I, SI Figure S11). Together, these results confirm that the altered predator-prey dynamics on chitin can be largely explained by a substantial decrease in VPS production.

## Discussion

Pathogenic *V. cholerae*’s life cycle involves shifts between the human host and the marine water column (3). In both host digestive tracts and marine conditions, *V. cholerae* forms biofilms to sequester space and growth resources, and to survive contact with a myriad of external threats including elements of the host immune system, bacteriophages, predators, and competing bacterial strains and species (2, 16, 23, 37). While *V. cholerae* is capable of forming pilus-dependent aggregates, it is generally thought to be VPS and related matrix proteins that contribute to robust *V. cholerae* biofilm formation (8, 33, 37, 41–45, 47, 50). How different growth conditions may alter the strength of matrix production, and how these changes cascade to changes in ecological interaction with other microbes, remain fruitful areas of work.

Here, we used the predatory bacterium *B. bacteriovorus* in conjunction with *V. cholerae* biofilm formation on chitin and glass substrates to explore how different environmental contexts influence *V. cholerae* biofilm architecture and predator-prey ecology (4, 31). We first showed that biofilm growth on chitin increases *V. cholerae*’s susceptibility to predation by *B. bacteriovorus*, while simultaneously decreasing the abundance of *B. bacteriovorus* from the system. We next used matrix deletion mutants cultivated on glass to recapitulate these key observations from the chitin experiments, and we demonstrated that VPS and matrix protein production is substantially reduced on chitin surfaces relative to glass substrate. Furthermore, the *V. cholerae* Toxin Co-regulated Pilus appears to play an under-appreciated role for biofilm integrity on chitin surfaces, and a more modest role in biofilm on glass surfaces. Finally, we showed that if VPS and related matrix protein production are induced on glass via genetic modification, biofilms regain key features of predator-prey interactions and biofilm architecture normally seen during glass cultivation.

Our finding that chitin-grown biofilms are more susceptible to predation than VPS-dependent biofilms grown on glass is suggestive of an important tradeoff that *V. cholerae* likely contends with in the marine environment. Pieces of chitin detritus offer a finite source of carbon and nitrogen, and producing large and densely-packed biofilms, such as those made by *vpvC^W240R^*, can offer competitive benefits and protection from exogenous threats. However biofilms produced by matrix hypersecretion have been shown to incur a cost in terms of dispersal and downstream colonization ability (51, 52). This suggests a tradeoff that could provide an evolutionary rationale for the existence *V. cholerae* variants from the marine environment that differ in their propensity to produce VPS and matrix proteins (49).

Additionally, the finding that VPS, RbmA, Bap1, and RbmC are utilized to a lesser degree on chitin relative to GlcNAc grown biofilms corresponds with prior reports of the Toxin Coregulated Pilus (TCP) and the Chitin regulated Competence Pilus (ChiRP) being required for biofilm formation on chitin (41, 46, 47). However, understanding the precise mechanisms underlying *V. cholerae* biofilm formation on chitin, and why they evolved a mechanistically distinct way of growing on chitin versus other surfaces, remain interesting and important areas for future work on the fundamental mechanisms of *V. cholerae* biofilm growth and the evolutionary dynamics of different strains of *V. cholerae* that switch between growing in hosts versus those that live exclusively in the marine environment.

Lastly, another interesting finding of our work is that *B. bacteriovorus* abundance on and within *V. cholerae* biofilms is dependent on production of VPS. The observation that VPS-dependent matrix both contributes to prey protection from predator exposure in the biofilm interior while also retaining some *B. bacteriovorus* cells in place suggests the possibility that biofilm architecture can stabilize predator-prey dynamics. This type of predator-prey interaction pattern may in turn contribute to the observed trophic diversity of naturally occurring biofilms (11, 53–57). While the mechanism by which VPS-dependent biofilm architecture affects *B. bacteriovorus* abundance in the biofilm interior is unknown, we speculate that VPS and matrix protein production lead to biofilm architectures that reduce overall prey accessibility for *B. bacteriovorus*, while also permitting some degree of entanglement that holds the predatory species in areas of the biofilm where matrix maturation has not yet occurred completely. Understanding the molecular mechanisms of this proposed interaction will be key to a fuller understanding of how biofilm matrix physiology influences microbial predator-prey dynamics more broadly.

## Materials and Methods

### Bacterial strains

The *V. cholerae* strains used in this study were derived from serotype O1 El Tor strain N1691 (58). The *B. bacteriovorus* strain was derived from 109J (30). All mutant derivatives were generated here and previously using standard allelic exchange methods with counter-selection for scarless deletion or integration of reporter constructs or point mutations (2, 5, 16, 23, 59). Cultures were grown overnight in lysogeny broth (LB, Miller) prior to inoculation into microfluidic devices. Biofilm cultures were grown in M9 minimal media, supplemented with 2 mM MgSO_4_, 100 μM CaCl_2_, and MEM vitamins, with either chitin flakes, Chitobiose ((GlcNAc)_2_) at 0.5%, or GlcNAc at 0.5% as the sole carbon source.

### Microfluidic flow device assembly

Microfluidic chambers used for biofilm culture were made with polydimethylsiloxane (PDMS) using standard soft lithography techniques (60, 61). PDMS was mixed and then cured on molds of chamber sets, punched with holes for inlet and outlet tubing, and bonded to #1.5 22 mm by 60 mm glass coverslips via plasma cleaning. The microfluidic chambers (7620 µm in length, 1500 µm in width and 80 µm in height) were designed to hold pieces of chitin flake in place using posts arranged in a ‘V’ shape, with the open end facing upstream into the flow path (SI Figure S1) (2). Prior to inoculation with bacteria, sterilized chitin flakes (Sigma) were loaded into the chambers, which were inspected to confirm that pieces of chitin were immobilized by the V-traps for subsequent *V. cholerae* colonization. Constant flow was applied via Harvard Apparatus Pico Plus syringe pumps loaded with 1 mL Brandzig plastic syringes. Syringes had 27-gauge needles affixed to them and fitted with #30 Cole Parmer PTFE tubing with an inner diameter of 0.3 mm. Tubing from the syringes was led to the inlets of each chamber, and outlet tubing was fed to a petri plate to collect waste or to Eppendorf tubes in instances where measurements were taken from the liquid exiting the chambers.

### Biofilm culture conditions

Strains were grown overnight in LB medium at 37° Celsius and shaken at 250 rpm. Overnight cultures were resuspended in M9 minimal media to an O.D._600_ of ∼4. For biofilms grown on chitin, molecular grade chitin flakes were incubated with ethanol for 1 min prior to being washed 5x with PBS and then resuspended in M9 with no carbon source. These chitin flakes were then flushed into microfluidic chambers to immobilize them in the V-trap sections of the devices (SI Figure S1) (2). For biofilms grown on glass with soluble carbon sources, the same microfluidic devices were used, but the disaccharide (GlcNAc)_2_ or the monosaccharide GlcNAc was added to the M9 media at 0.5%. While the exact ratio of soluble sugar produced by chitin degradation remains unclear, (GlcNAc)_2_ and GlcNAc are likely end-products of *Vibrio* chitinases (4). After loading the chambers with chitin flakes, 10 μL of inoculum was transferred into the chambers and allowed to stay in place without flow for 1 h to permit bacterial colonization of the chitin particles. After this colonization step, M9 minimal media was introduced to the chambers at a rate of 0.1 µl/min (corresponding to an average flow velocity of ∼ 14 µm/s) at room temperature (∼22° Celsius).

### Biofilm predation assay

After 72 h of biofilm growth, the inlet media was swapped for 2 h to either (1) M9 media containing *B. bacteriovorus* resuspended in M9 media at a concentration of 10^10^ plaque forming units (PFUs) per mL; or (2) for control chambers, the same inlet swap procedure was performed, but to a media supply that did not contain any *B. bacteriovorus*. Following the 2 h introduction of predator bacteria or sterile control, the media inlet was then swapped back to the original sterile M9 media, and flow was paused for 30 min to allow for adherence of the invading *B. bacteriovorus*. Flow of sterile media was then resumed for the duration of the experiment. Imaging was performed at 4 h post invasion and at 24 h intervals thereafter. The biovolume of strains, as a proxy for their population size, was measured by imaging z-stacks of multiple 212×212 µm fields of view within each replicate chamber.

### Predation localization

To quantify the probability of predation with respect to biofilm local cell packing, the biofilm predation assay was repeated as previously described, but imaging was performed 2 h after predator introduction. Given that the latent period of *B. bacteriovorus* is as short as 3-4 h, this 2 h time point ensured that images were collected prior to *V. cholerae* predation-driven cell lysis (62–64). We then aggregated our image data from the three treatments such that there was a uniform distribution of local cell packing values before calculating the probability of predation with respect to local biofilm structure.

### Percolation model of predation

To model the distribution of predation events with respect to local cell packing, we used a directed percolation model (65). A 3-dimensional grid of size *L*x*L*x*L* was randomly populated with site occupation probabilities ranging from 0 to 1. For each site occupation probability, *N* random grid samples were generated. A depth-first search for a connected path from outside to the center through unoccupied grid points was performed on each grid. If a path from outside to the center of the 3-dimensional grid was found, that simulation was counted as a successful predation event. If a path was not found, that simulation was counted as a failed predation event. For a graphical illustration in two dimensions detailing how this model relates to our experimental data see SI Figure S3. SI Table S1 provides a list of parameters and units. The dimensions of the 3-D grids used for this model were matched to the dimensions used to calculate local biofilm bacterial volume fractions (cell packing values) used for analysis of the experimental data.

### Predator-prey model

To model predator-prey dynamics, we used a classical Lotka-Volterra model (66). We monitor a prey species, *N* (μm^3^), that grows in a density dependent manner: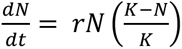, where *r* is the maximal growth rate (d^−1^), and *K* is the carrying capacity (μm^3^). We then introduce a predator, *P* (μm^3^), that has a loss rate *d* (d^−1^) and attacks the prey species at rate *α* (μm^−3^ d^−1^), generating *B. bacteriovorus* cells that are actively digesting infected prey, termed bdelloplasts (*B*, um^3^). Bdelloplasts mature into new predators with conversion efficiency *b* (unitless) and rate *k_p_* (d^−1^), giving the set of equations:

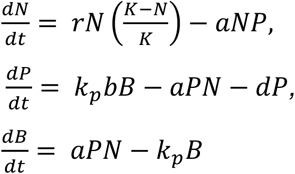

To simulate the effect of biofilm local cell packing blocking predator access, we added an additional interaction term, *v*, giving:

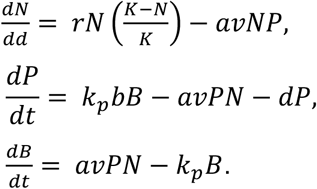

For a list of parameters see SI Table S3.

### Matrix component staining assay

RbmA, Bap1, and RbmC were engineered to carry a C-terminal 3xFLAG epitope as previously described (16, 25, 34). A Cy3-conjugated anti-FLAG monoclonal antibody was then introduced into the culture medium for the duration of the experiment at 1 µg/1 mL To stain VPS, Alexa fluor 488 was fused to Bap1 and purified protein was spiked into the chambers at a concentration of 1 μM and allowed to incubate without flow for 0.5 h (36). After 0.5 h, flow was resumed to wash out unbound stain. Biofilms were grown as described in the biofilm culture conditions section. Stain signal was quantified within a 0.75 μm shell of *V. cholerae* biovolume.

### Biofilm competition assay

Competing strains were inoculated into microfluidic devices at an O.D._600_ of ∼4 at a range of initial ratios. Starting ratios were then experimentally confirmed 1 h post inoculation via confocal microscopy. At 5 d post inoculation the relative abundances of the competitors were again measured via confocal microscopy. Biofilms were maintained as described in the biofilm culture conditions section.

### Fluorescence microscopy

All imaging was performed using a Zeiss 880 line-scanning confocal microscope with a 40x/1.2 N.A. water objective. The GFP protein expressed constitutively by *B. bacteriovorus* was excited with a 488 nm laser line. The mKO-κ protein that *V. cholerae* expresses constitutively was excited with a 543 nm laser line. The mKate2 protein that *V. cholerae* expresses constitutively was excited with a 594 nm laser line. The Cy3 Anti-FLAG was excited with a 543 nm laser line. The Bap1-Alexa488 fusion was excited with a 488-nm laser line. The chitin, which is auto-fluorescent, was excited with a 405 nm laser line. All representative images were processed by constrained iterative deconvolution in ZEN Blue.

### Image analysis

CZI image files were converted to TIFF format prior to being loaded into either a custom Python script or BiofilmQ, which was run using MATLAB (67). Thresholding was done using Otsu’s method with a manual sensitivity adjustment (68). Cube side lengths were set to 2.3 µm, giving cubes that can maximally hold ∼12 individual cells. For spatially resolved analyses, the thresholded 3-dimentional bacterial populations were segmented into a cubic grid for quantification of parameters at a pseudo-cell resolution. Calculated parameters were then exported from BiofilmQ as .mat files and loaded into Python, where SciPy, seaborn, Pandas, NumPy, and Matplotlib were used for running statistical tests and graphing (69–76). For calculated cube-based parameters that require a range, we chose 6 µm as the range over which to measure (see main text, SI Figure S3, and *Wucher* et al. 2021) (16). To account for cubes containing varying amounts of biovolume, cube-based histograms are weighted by the volume contained withing each cube. A custom python script was used to correlate the VPS and RbmA stains. Operation of BiofilmQ and its array of analytical methods is described in extensive detail in *Hartmann* et al. (77). All custom scripts are publicly available on <u>GitHub</u>.

### Replication and statistics

At least three biological replicates, each defined as the average of technical replicate image stacks within a single flow chamber, were collected for each experiment. For biovolume measurements, multiple regions of 212×212 µm were sampled as technical replicates within a chamber and averaged for each biological replicate. Z-stacks for all experiments were collected at a height of 15 μm due to aberration caused by the chitin at greater imaging depths. For all time-resolved data, the error bars correspond to one standard error. For box and whisker plots, the orange bar denotes the median, the box denotes Q1 and Q3, and the whiskers denote Q1 and Q3 + 1.5 times the interquartile range. The Python scripts and corresponding datasheets used to run models, generate figures, and perform statistical analysis are publicly available on [GitHub](https://github.com/nadellinsilico/Bdello_Chitin_Predation_2025).

## Acknowledgments

We are grateful to the members of the Nadell group and the microbiology communities at Dartmouth for feedback on the project over the course of its development. The study was supported by the Simons Foundation Award No. 826672 and NIGMS grant 1R35GM151158-01 to C.D.N.

## SI Figures

**SI Figure S1:**
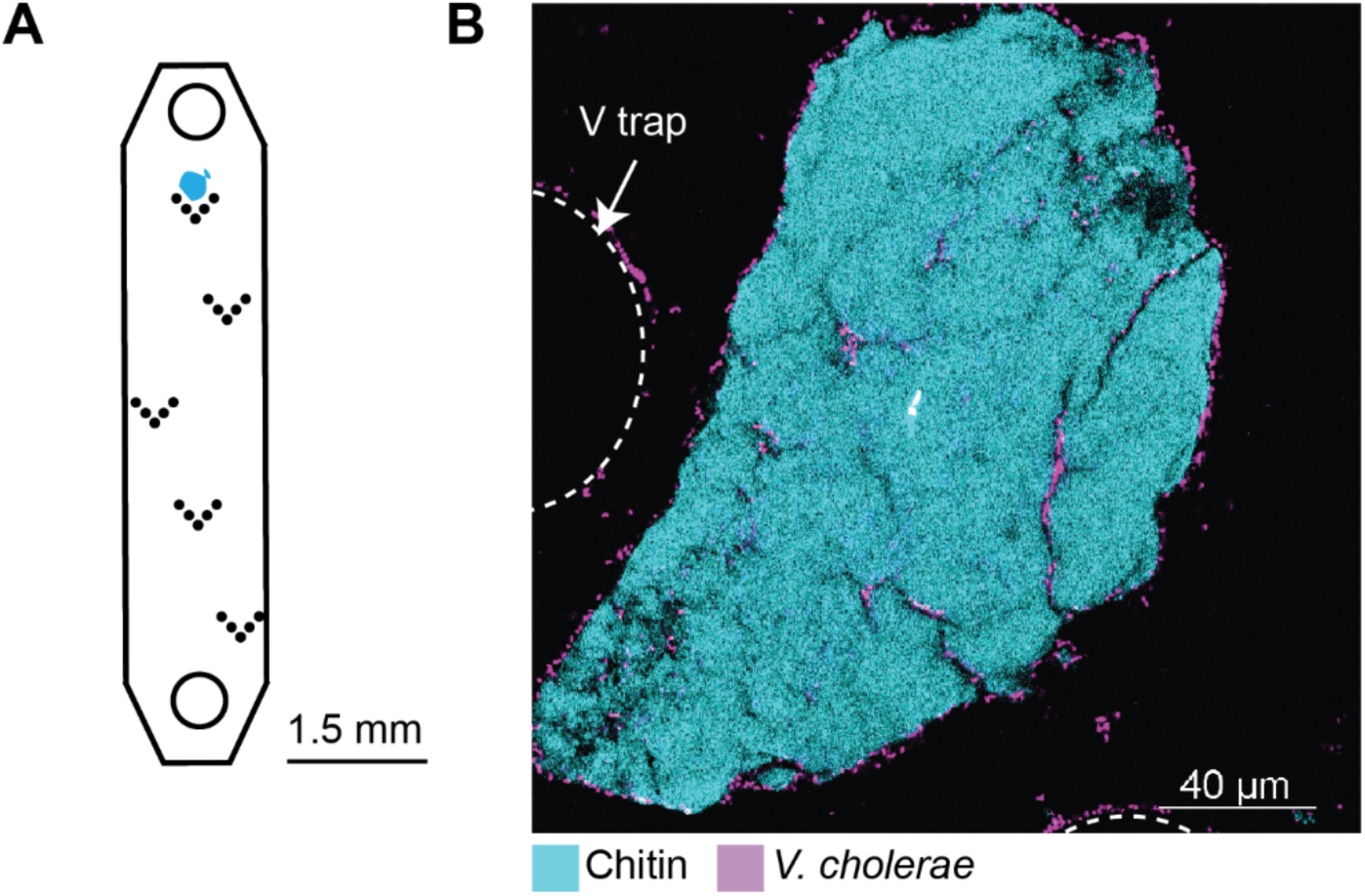
V trap microfluidic device. **(A)** Cartoon drawing showing a V trap microfluidic device chamber. Chamber drawn to scale. **(B)** Representative image of a *V. cholerae* colonized chitin flake wedged between two columns of a v trap. Chitin is shown in cyan, *V. cholerae* is shown in purple, and the v traps are shown as white, dashed lines

**SI Figure S2:**
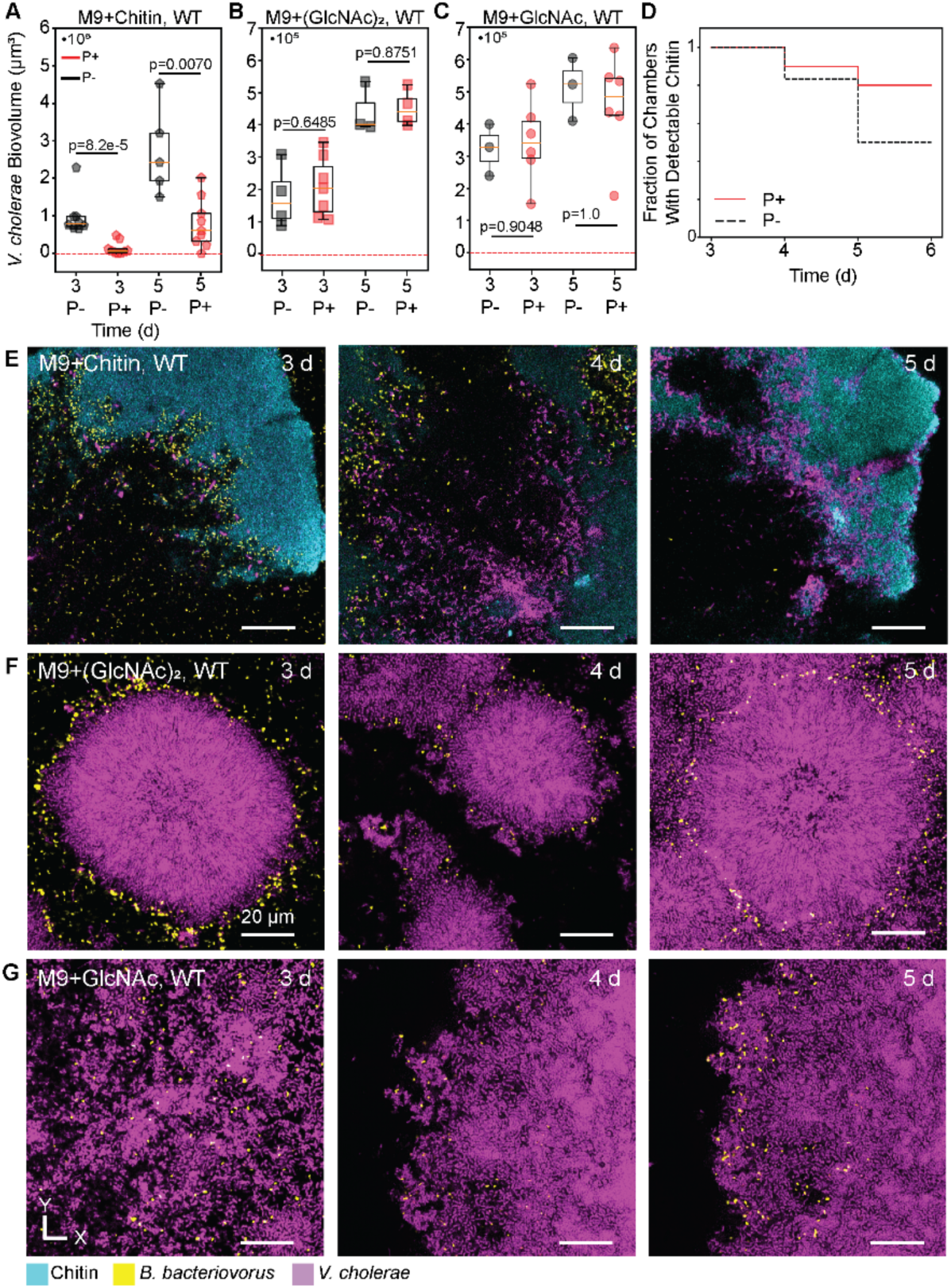
*V. cholerae* biofilms grown on chitin have increased susceptibility to predation relative to biofilms grown in GlcNAc or (GlcNAc)_2_. **(A)** Box and whisker plot comparing predation to no predation chitin biofilms at 3 d (∼4 h post introduction of predator) and 5 d, showing a significant decrease in *V. cholerae* biofilm abundance (Mann-Whitney *U* test, n=5-9). **(B)** Box and whisker plot comparing predation to no predation (GlcNAc)_2_ biofilms at 3 d (∼4 h post introduction of predator) and 5 d showing no significant difference in *V. cholerae* biofilm abundance (Mann-Whitney *U* test, n=3-7). **(C)** Box and whisker plot comparing predation to no predation GlcNAc biofilms at 3 d (∼4 h post introduction of predator) and 5 d showing no significant decrease in *V. cholerae* biofilm abundance (Mann-Whitney *U* test, n=3-6). **(D)** Fraction of microfluidic chambers with detectable chitin plotted against time for the *V. cholerae* chitin formed biofilms that were invaded by *B. bacteriovorus*, shown as the red line, and biofilms that were not invaded, shown as the dashed black line. **(E)** Representative images of *V. cholerae* biofilms being predated upon by *B. bacteriovorus* when grown with chitin as the sole carbon source. *V. cholerae* is shown in purple, *B. bacteriovorus* is shown in yellow, and chitin is shown in cyan. **(F)** Representative images of *V. cholerae* biofilms being predated upon by *B. bacteriovorus* when grown with (GlcNAc)_2_ as the sole carbon source. *V. cholerae* is shown in purple, *B. bacteriovorus* is shown in yellow, and chitin is shown in cyan. **(G)** Representative images of *V. cholerae* biofilms being predated upon by *B. bacteriovorus* when grown with GlcNAc as the sole carbon source. *V. cholerae* is shown in purple, *B. bacteriovorus* is shown in yellow, and chitin is shown in cyan.

**SI Figure S3:**
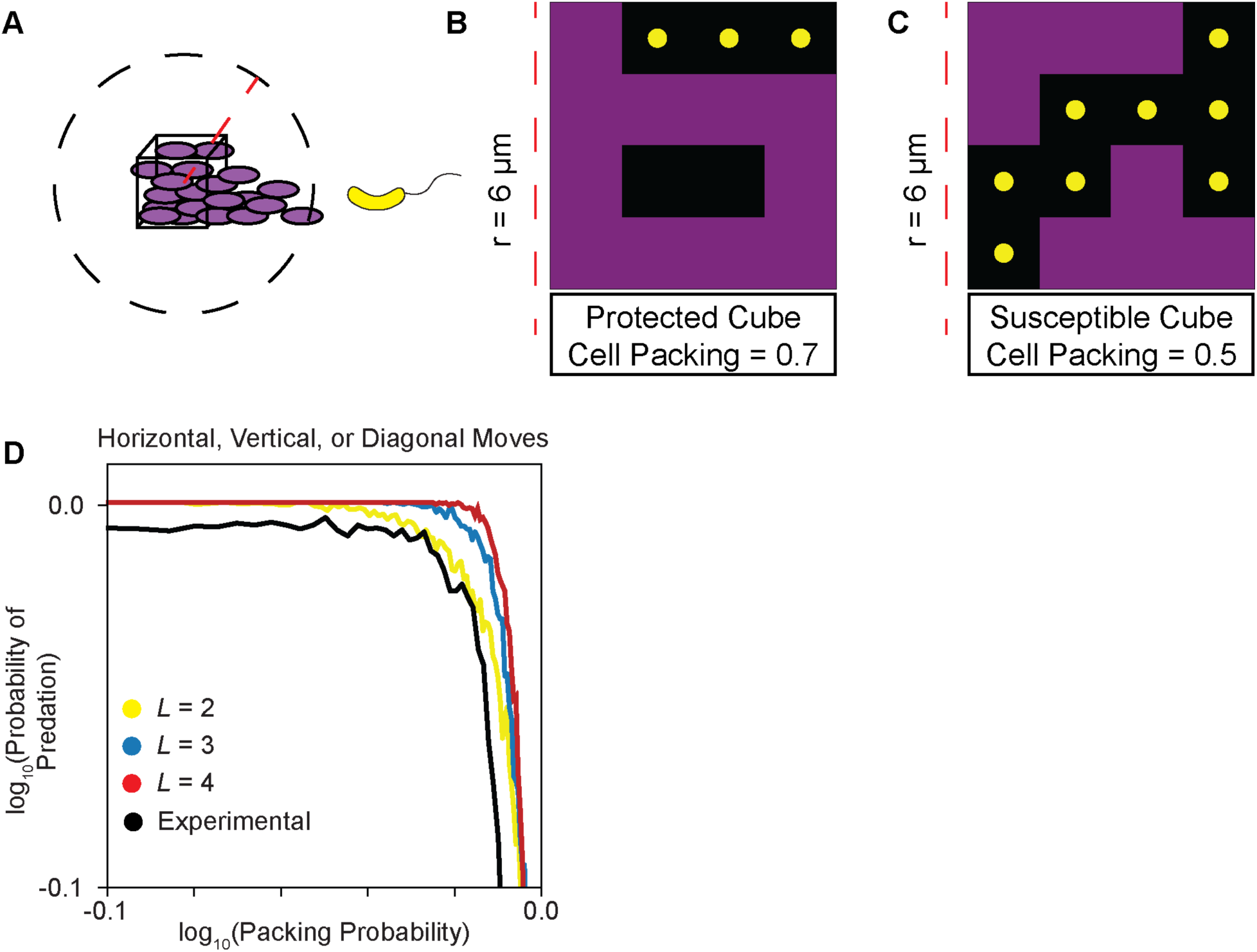
Percolation model. **(A)** Diagram detailing how BiofilmQ calculates local cell packing. (**B)** Representative 2D simulation of percolation with a cell packing fraction of 0.7 and grid size of 5. **(C)** Representative 2D simulation of percolation with a cell packing fraction of 0.5 and grid size of 5. *V. cholerae* is shown in purple, *B. bacteriovorus* is shown in yellow, the BiofilmQ radius parameter is shown as a dashed red line, the volume over which BiofilmQ calculates local cell packing fraction is shown as a black dashed line, and the BiofilmQ cube-object is shown as a cube in A and a rectangle in B and C. **(D)** Parameter sweep of the 3D percolation model where the grid size, *L*, is varied from 2 to 4 and diagonal moves are allowed.

**SI Figure S4:**
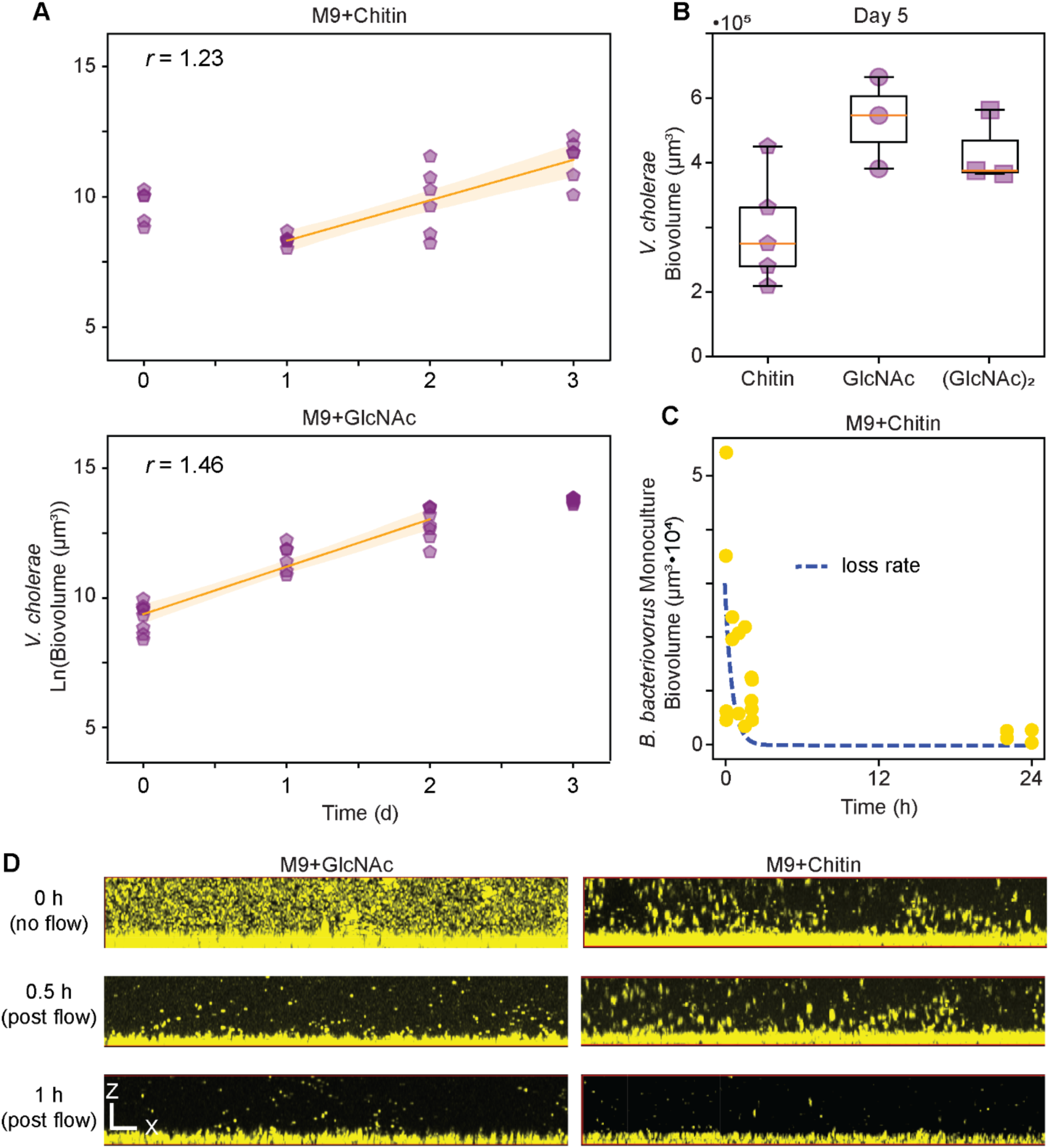
Monoculture data used to parameterize the Lotka-Volterra model. **(A)** Plot of natural log of biovolume against time for WT *V. cholerae* grown in M9 with chitin flakes or M9 with GlcNAc as the sole carbon source. Solid orange line represents the line of best fit used to experimentally determine the intrinsic rate of increase, *r*. **(B)** Box and whisker plot of WT *V. cholerae* biofilm volume at day 5, used to determine the carrying capacity, *K*. **(C)** Population dynamics of *B. bacteriovorus* over 24 h in microfluidic devices not containing prey biofilm. The dashed blue line is the microfluidic device dilution rate. **(D)** Representative x-z maximal intensity projection series of *B. bacteriovorus* monoculture in M9+Chitin (right) and in M9 with GlcNAc (left). *B. bacteriovorus* is shown in yellow.

**SI Figure S5:**
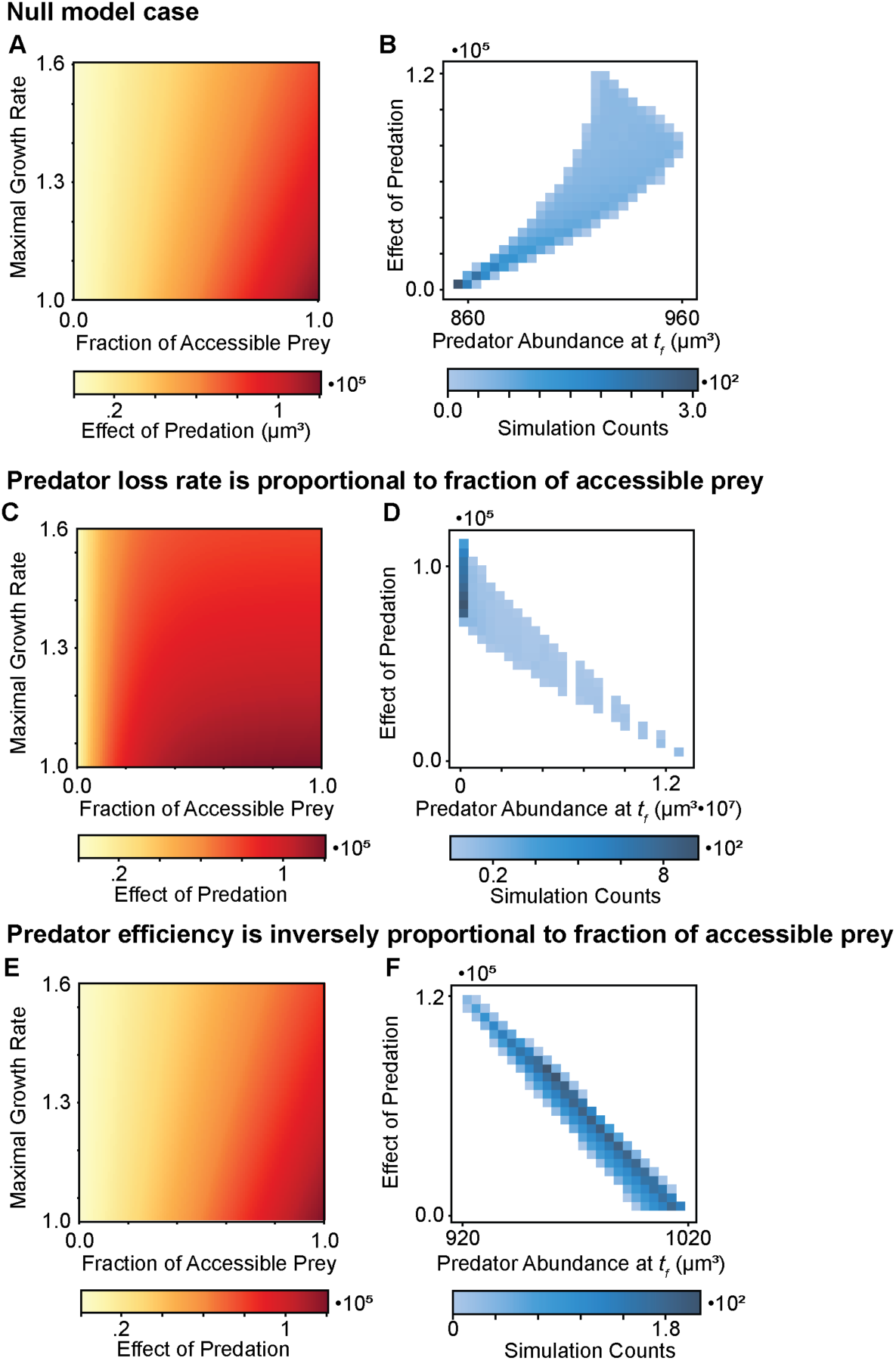
Response of predator abundance as model assumptions are altered. **(A)** Plot of the predator’s effect on its prey as the fraction of accessible prey and the maximal growth rate are varied. Replot of the simulation results from Figure 1I. **(B)** 2D histogram of predator abundance at the simulation end (6 d) against effect of predation for all model runs showing that an increase in accessible prey correlates positively with predator abundance. **(C)** Plot of the predator’s effect on its prey as the fraction of accessible prey and the maximal growth rate are varied in the model case where the predator’s loss rate is modified proportionally to the fraction of accessible prey. **(D)** 2D histogram of predator abundance at the simulation end (6 d) against effect of predation for all model runs showing that an increase in accessible prey correlates positively with predator abundance. **(E)** Plot of the predator’s effect on its prey as the fraction of accessible prey and the maximal growth rate are varied in the model case where the predator’s efficiency is modified inversely to the fraction of accessible prey. **(F)** 2D histogram of predator abundance at the simulation end (6 d) against effect of predation for all model runs showing that an increase in accessible prey correlates positively with predator abundance.

**SI Figure S6:**
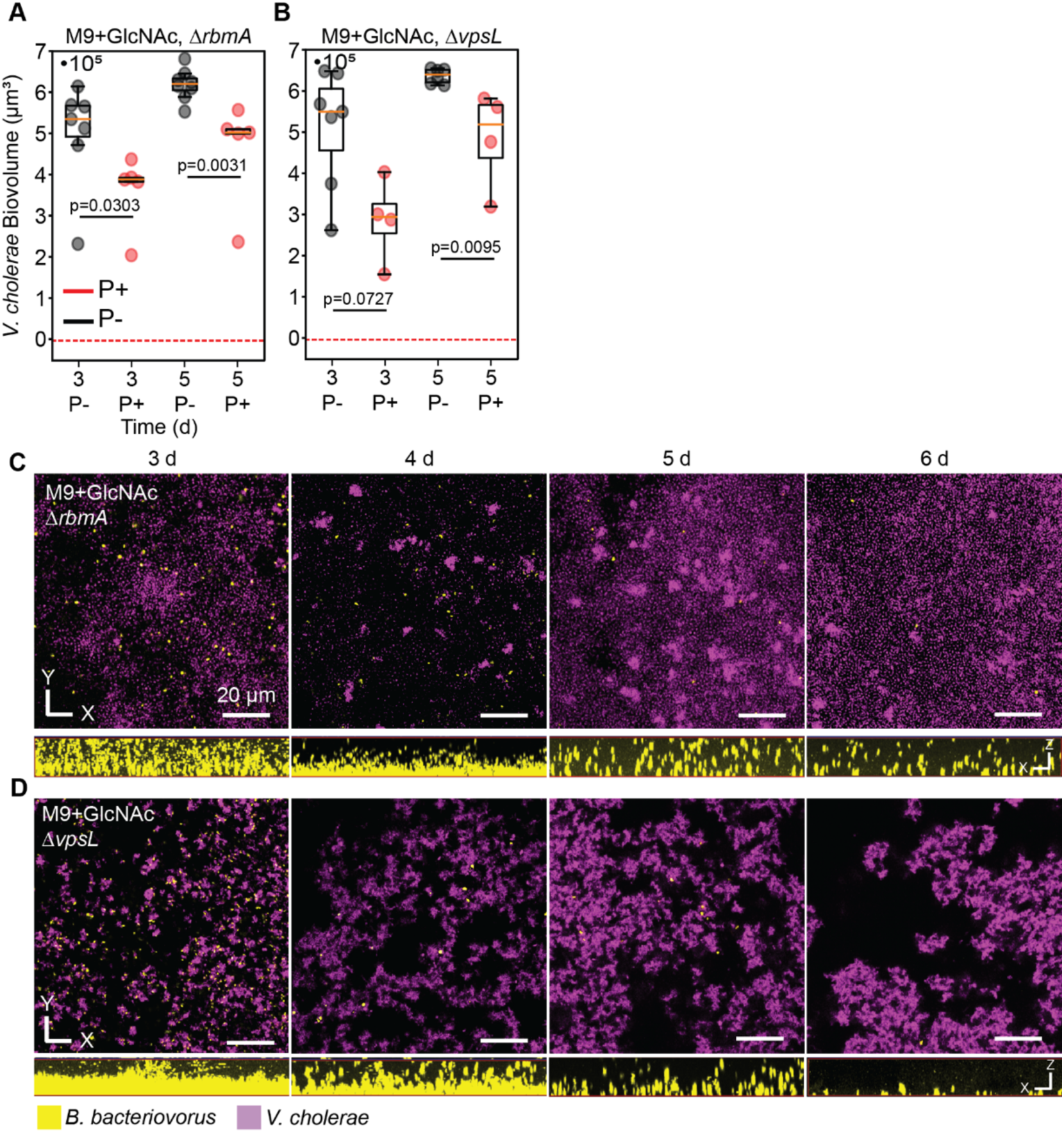
Dense local cell packing is necessary for biofilm level protection against *B. bacteriovorus,* and VPS secretion is necessary for *B. bacteriovorus* maintenance. **(A)** Box and whisker plot comparing predation to no predation *ΔrbmA* biofilms grown in GlcNAc at 3 d (∼4 h post introduction of predator) and 5 d showing a significant decrease in *V. cholerae* biofilm abundance (Mann-Whitney *U* test, n=4-7). **(B)** Box and whisker plot comparing predation to no predation *ΔvpsL* biofilms grown in GlcNAc at 3 d (∼4 h post introduction of predator) and 5 d showing a significant difference in *V. cholerae* biofilm volume (Mann-Whitney *U* test, n=4-7). **(C)** Representative images of *V. cholerae ΔrbmA* biofilms being predated upon by *B. bacteriovorus* over time with x-z maximum intensity projections of *B. bacteriovorus* shown below. **(D)** Representative images of *V. cholerae ΔvpsL* biofilms being predated upon by *B. bacteriovorus* over time with x-z maximum intensity projections of *B. bacteriovorus* shown below. *V. cholerae* is shown in purple and *B. bacteriovorus* is shown in yellow.

**SI Figure S7:**
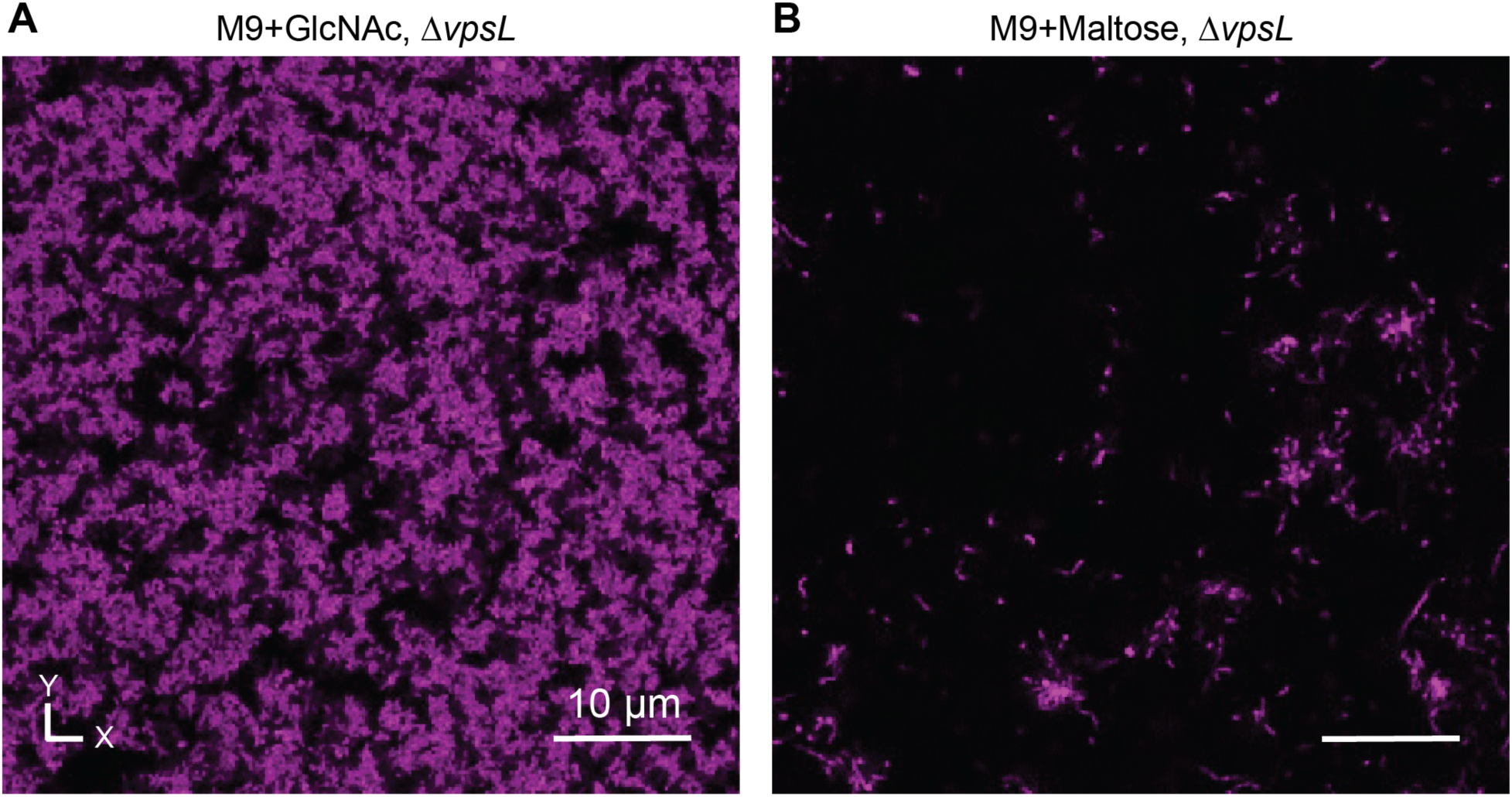
*V. cholerae ΔvpsL* forms disordered 3-dimensional biofilms dependent on media conditions. **(A)** *V. cholerae ΔvpsL* forms 3-dimensional biofilms when grown with GlcNAc as the sole carbon source. **(B)** *V. cholerae ΔvpsL* does not form 3-dimensional biofilms when grown with maltose as the sole carbon source. *V. cholerae ΔvpsL* is shown in purple.

**SI Figure S8:**
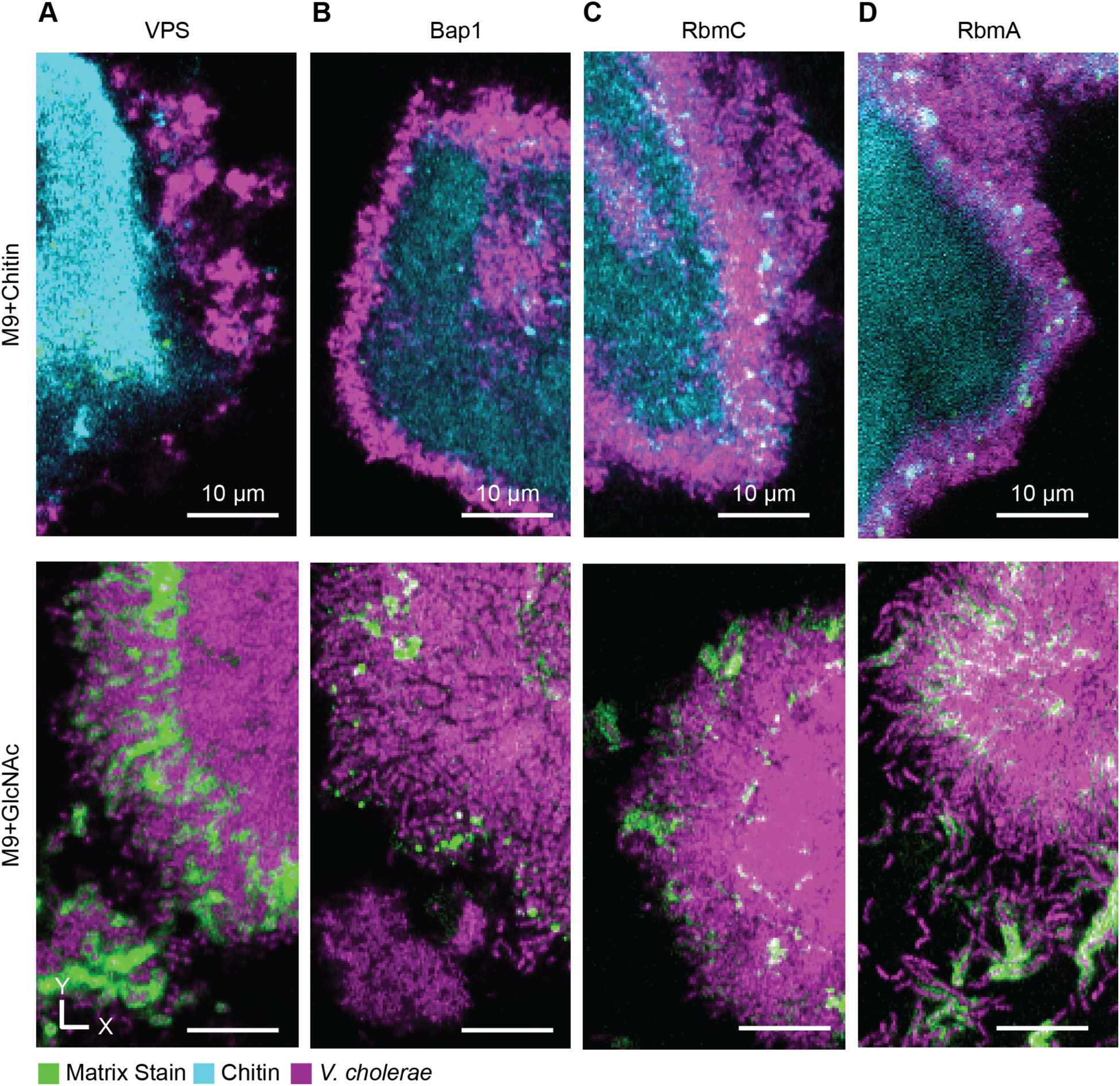
*V. cholerae* biofilms formed on chitin contain less *Vibrio* exopolysaccharide, RbmA, RbmC, and Bap1 relative to biofilms formed in GlcNAc. **(A)** Representative image of *V. cholerae* grown on chitin (top) stained for VPS alongside a representative image of *V. cholerae* grown in GlcNAc also stained for VPS (bottom). **(B)** Representative image of *V. cholerae* grown on chitin (top) stained for Bap1 alongside a representative image of *V. cholerae* grown in GlcNAc also stained for Bap1 (bottom). **(C)** Representative image of *V. cholerae* grown on chitin (top) stained for RbmC alongside a representative image of *V. cholerae* grown in GlcNAc also stained for RbmC (bottom). **(D)** Representative image of *V. cholerae* grown on chitin (top) stained for RbmA alongside a representative image of *V. cholerae* grown in GlcNAc also stained for RbmA (bottom). In all image panels matrix stain is shown in green, chitin is shown in cyan, and *V. cholerae* is shown in purple.

**SI Figure S9:**
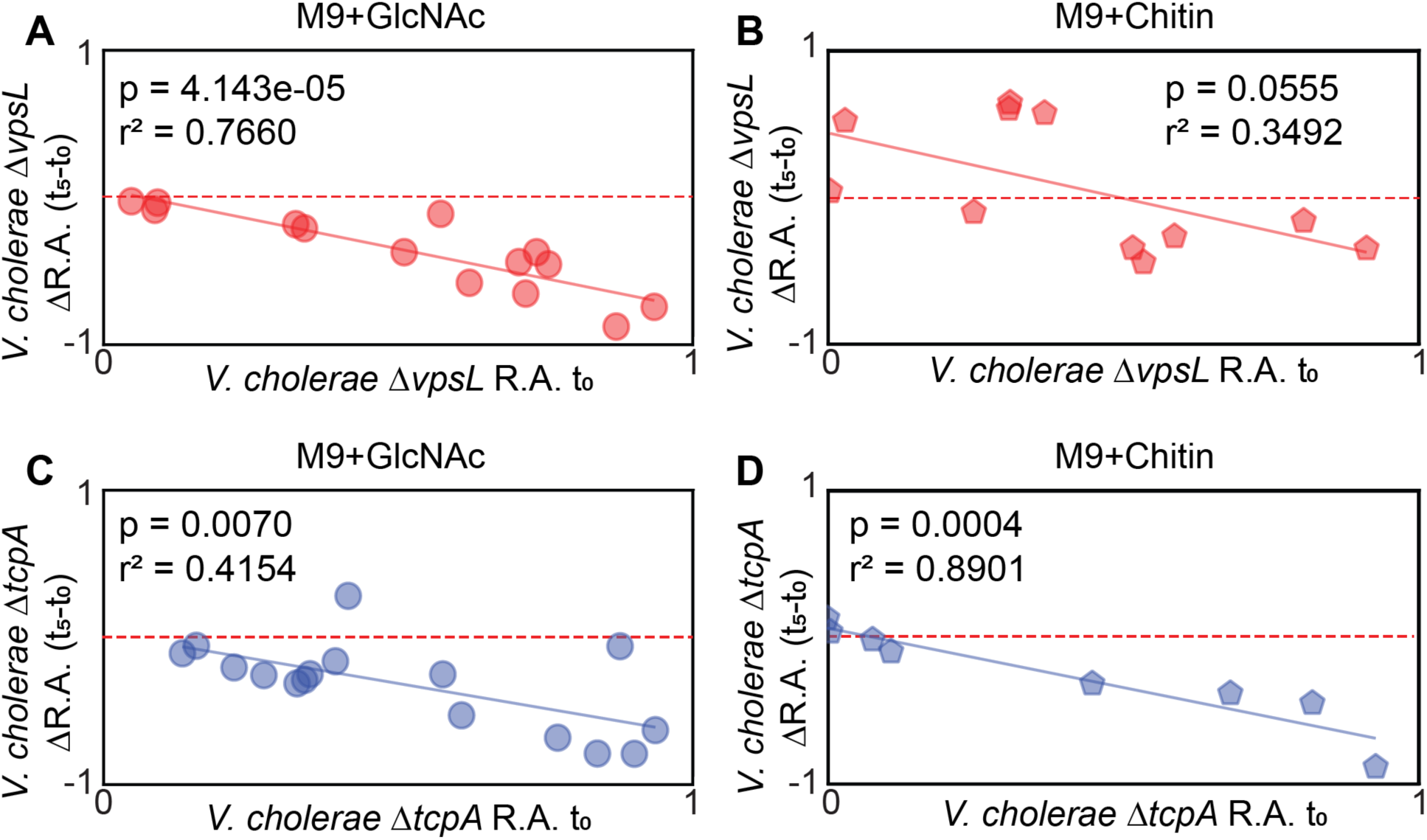
Growth substrate alters the competitive ability of WT against *ΔvpsL* and *ΔtcpA* **(A)** Plot of *V. cholerae ΔvpsL* initial relative abundance against the change in *V. cholerae ΔvpsL* relative abundance after five days of coculture with WT grown in M9 with GlcNAc as the sole carbon source. **(B)** Plot of *V. cholerae ΔvpsL* initial relative abundance against the change in *V. cholerae ΔvpsL* relative abundance after five days of coculture with WT grown in M9 with Chitin as the sole carbon source. **(C)** Plot of *V. cholerae ΔtcpA* initial relative abundance against the change in *V. cholerae ΔtcpA* relative abundance after five days of coculture with WT grown in M9 with GlcNAc as the sole carbon source. **(D)** Plot of *V. cholerae ΔtcpA* initial relative abundance against the change in *V. cholerae ΔtcpA* relative abundance after five days of coculture with WT grown in M9 with chitin as the sole carbon source. The red dashed line shows the zero-change line.

**SI Figure S10:**
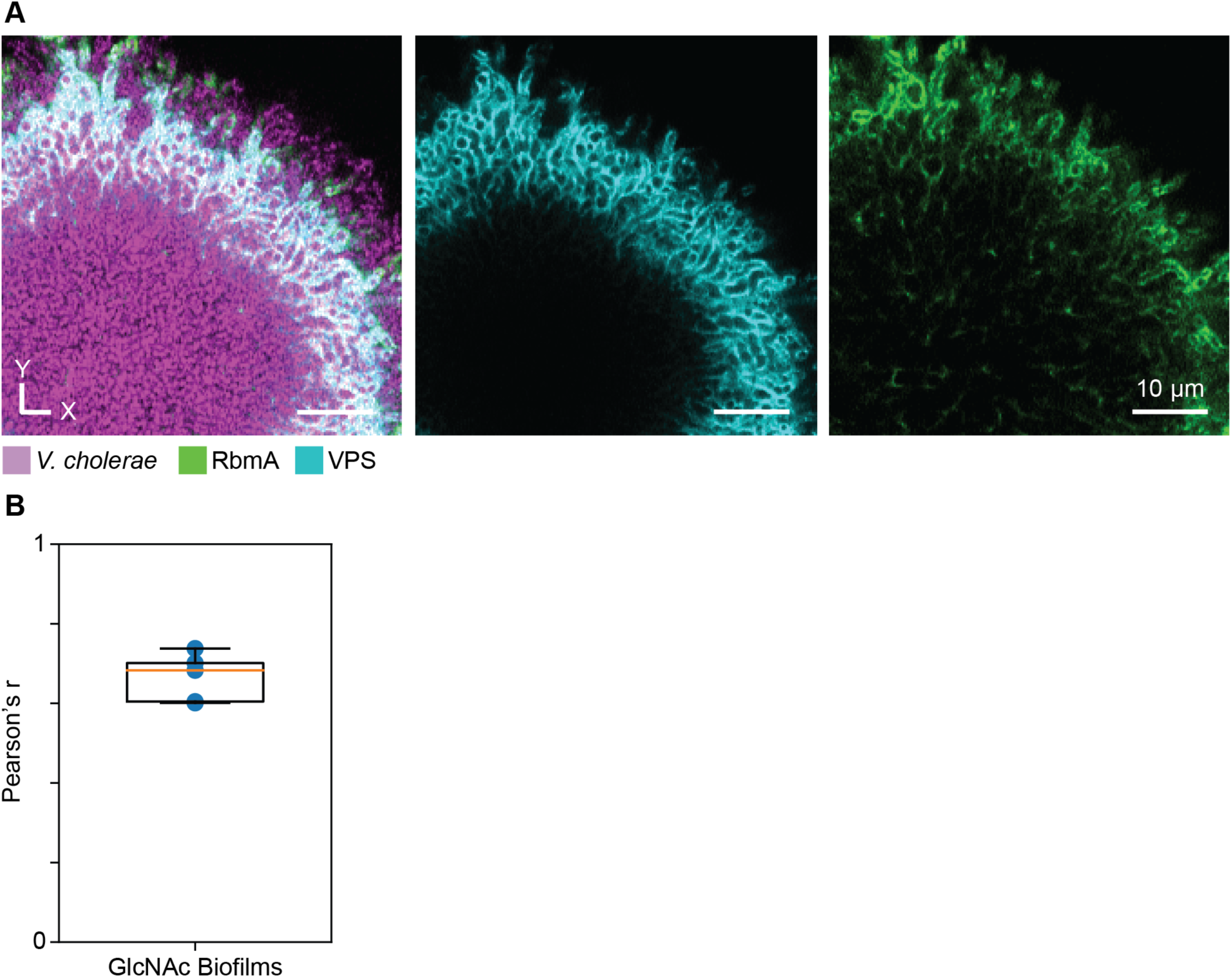
VPS stain is strongly correlated with RbmA stain. **(A)** Representative image of a V. cholerae biofilm grown in GlcNAc exposed continuously to RbmA for 2 d and then stained with a spike in of Bap1 bound to Alexa Fluor 488. *V. cholerae* is shown in purple, VPS is shown in cyan, and RbmA is shown in green. **(B)** Pearson’s correlation coefficient of VPS and RbmA stain signal (n = 5 images).

**SI Figure S11:**
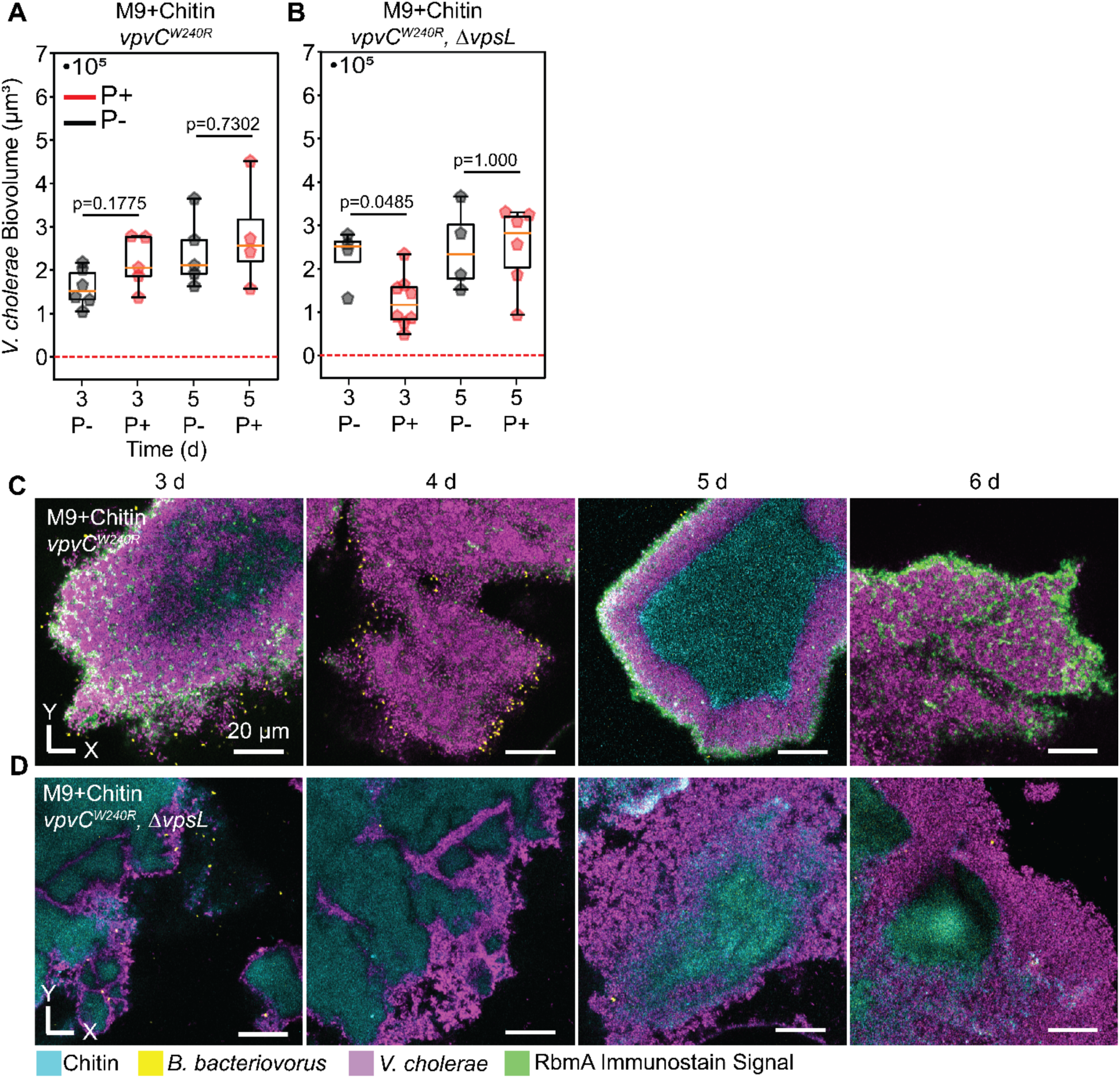
*V. cholerae vpvC^W240R^* biofilms are protected from predation in a VPS-dependent manner when grown on chitin. **(A)** Box and whisker plot comparing predation to no predation *vpvC^W240R^*biofilms grown on chitin at 3 d (∼4 h post introduction of predator) and 5 d showing no significant difference in *V. cholerae* biofilm volume (Mann-Whitney *U* test, n=4-6). **(B)** Box and whisker plot comparing predation to no predation *vpvC^W240R^ ΔvpsL* biofilms at 3 d (∼4 h post introduction of predator) and 5 d showing a significant difference (Mann-Whitney *U* test, n=4-6). **(C)** Representative images of *V. cholerae vpvC^W240R^* biofilms being predated upon by *B. bacteriovorus* over time. **(D)** Representative images of *V. cholerae vpvC^W240R^ ΔvpsL* biofilms being predated upon by *B. bacteriovorus* over time. *V. cholerae* is shown in purple, *B. bacteriovorus* is shown in yellow, chitin is shown in cyan, and RbmA immunostaining signal is shown in green.

## SI Tables

**SI Table S1:**
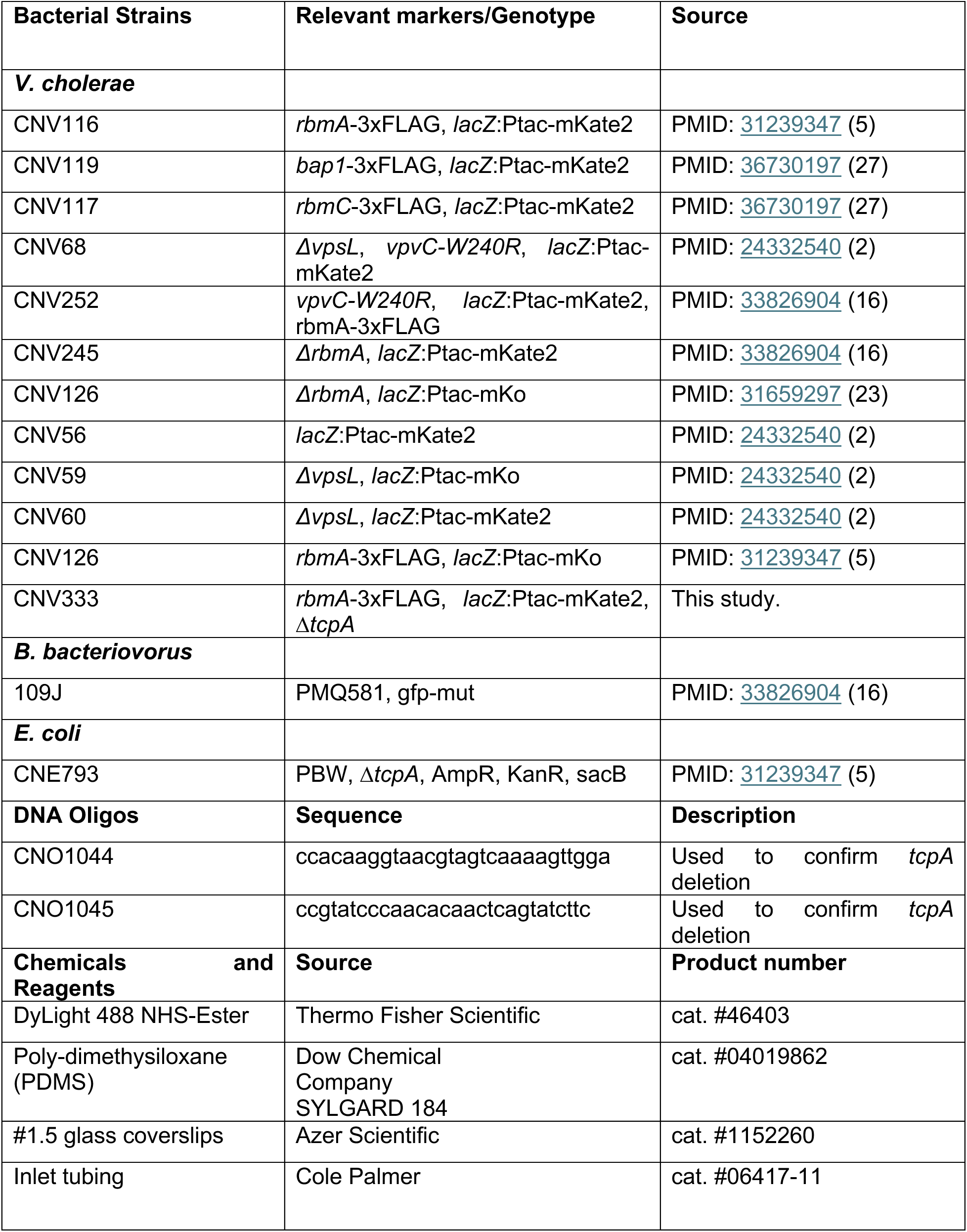
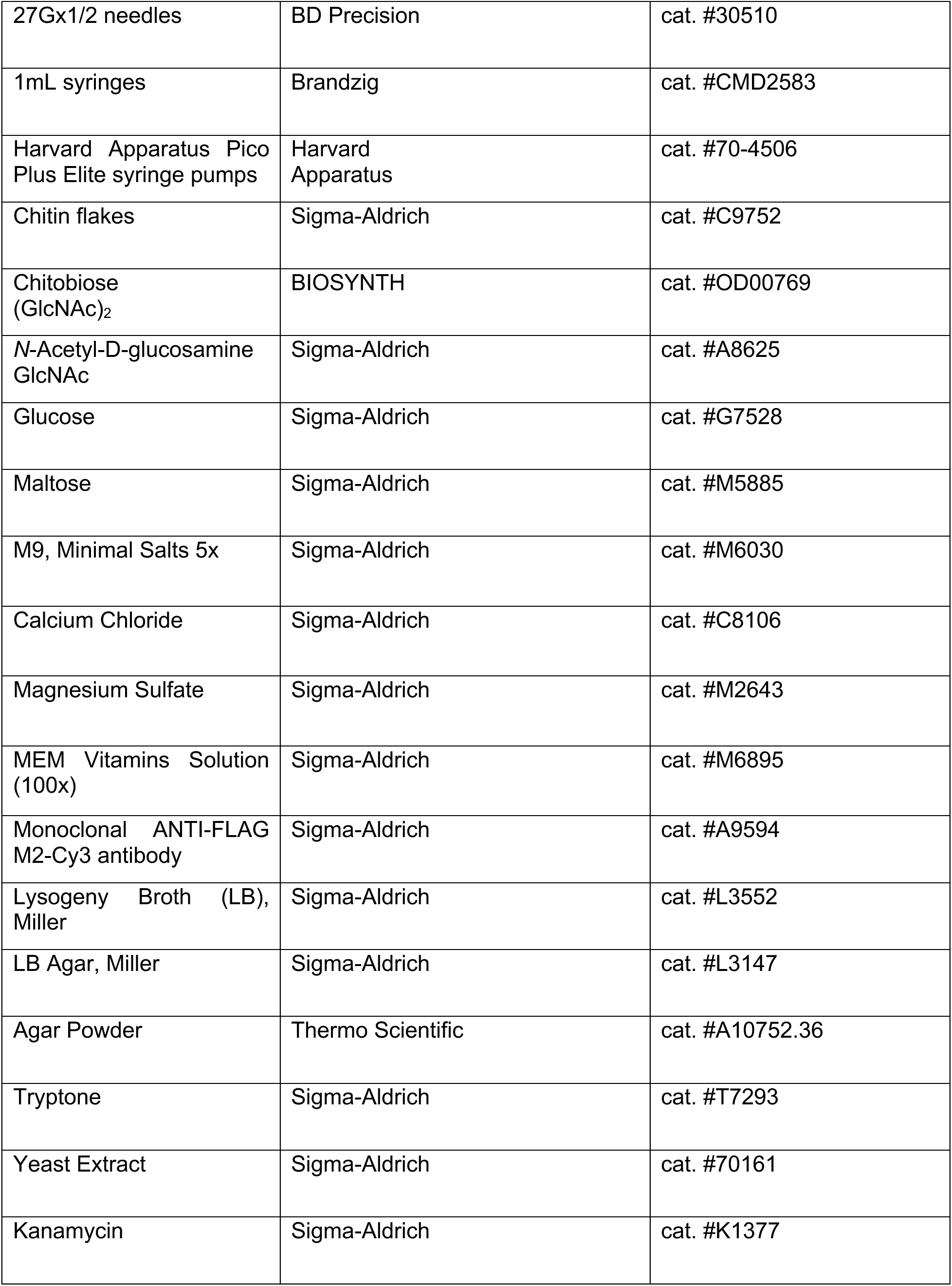
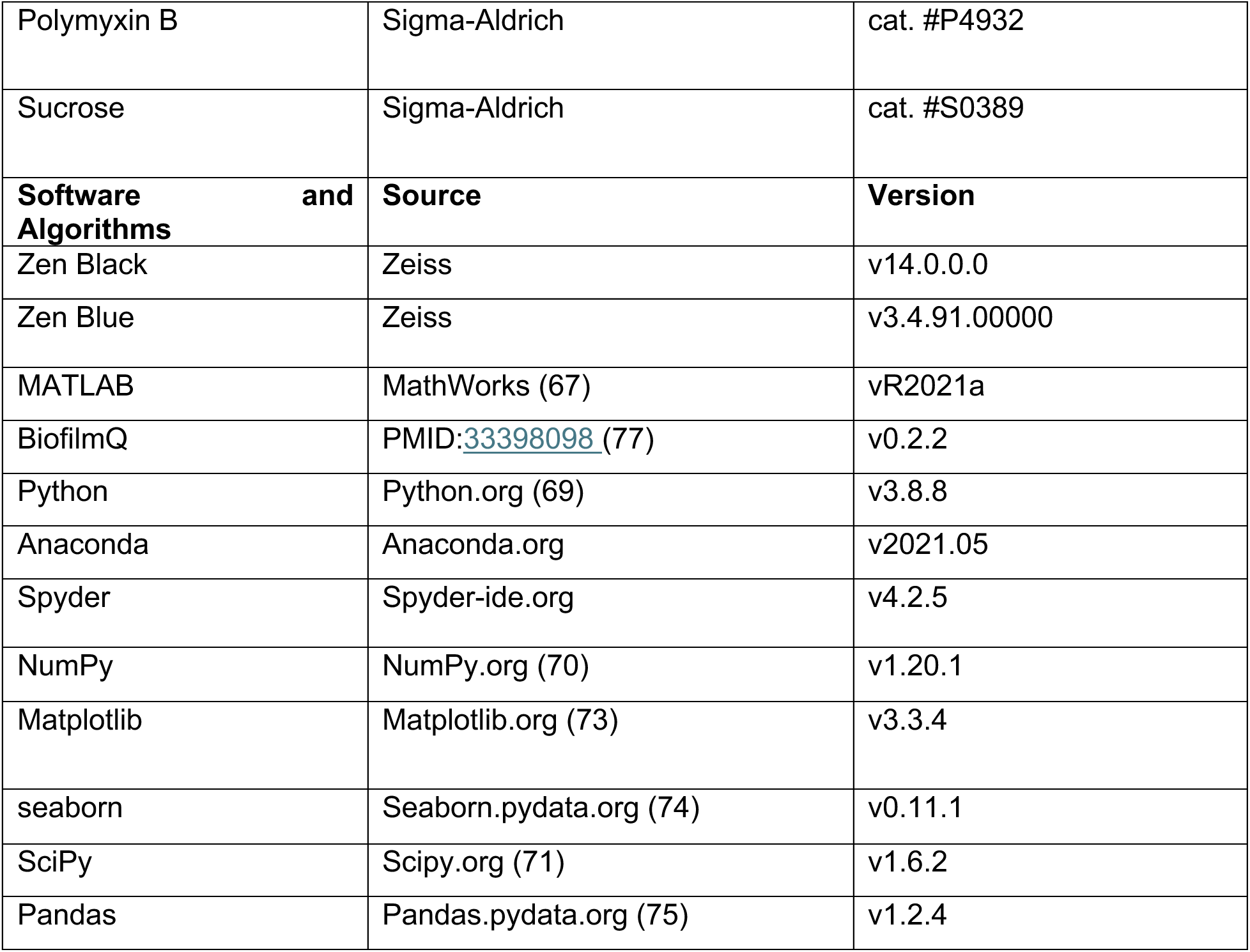
List of strains, materials, and software.

**SI Table S2:** List of percolation model parameters.

| Parameter | Symbol | Value | Unit |
| --- | --- | --- | --- |
| Grid size | $L$ | 2-4 | individual cells<br>(~0.5-2.5 $\mu\text{m}$) |
| Samples | $N$ | 200 | unitless |

**SI Table S3:** List of ODE model parameters.

| Parameter | Symbol | Value | Unit |
| --- | --- | --- | --- |
| *V. cholerae* maximal growth rate | $r$ | 0.5-1.5<br>(SI Figure S4) | $\text{d}^{-1}$ |
| *V. cholerae* carrying capacity | $K$ | $3 \cdot 10^5$-$5 \cdot 10^5$<br>(SI Figure S4) | $\mu\text{m}^3$ |
| *B. bacteriovorus* attack rate | $a$ | $5.52 \cdot 10^{-7}$ (64) | $\mu\text{m}^{-3} \text{d}^{-1}$ |
| *B. bacteriovorus* loss rate | $d$ | 1.7952<br>(SI Figure S4) | $\text{d}^{-1}$ |
| Bdelloplast maturation rate | $k_p$ | 2.616 (63) | $\text{d}^{-1}$ |
| Bdelloplast to predator conversion factor | $b$ | 0.438 (63) | unitless |
| Predator-prey interaction term | $v$ | 0-1<br>(Figure 1,<br>SI Figure S5) | unitless |

